# A Comprehensive Classification of Coronaviruses and Inferred Cross-Host Transmissions

**DOI:** 10.1101/2020.08.11.232520

**Authors:** Yiran Fu, Marco Pistolozzi, Xiaofeng Yang, Zhanglin Lin

**Affiliations:** School of Biology and Biological Engineering, South China University of Technology, 382 East Outer Loop Road, University Park, Guangzhou 510006, China

## Abstract

In this work, we present a unified and robust classification scheme for coronaviruses based on concatenated protein clusters. This subsequently allowed us to infer the apparent “horizontal gene transfer” events via reconciliation with the corresponding gene trees, which we argue can serve as a marker for cross-host transmissions. The cases of SARS-CoV, MERS-CoV, and SARS-CoV-2 are discussed. Our study provides a possible technical route to understand how coronaviruses evolve and are transmitted to humans.

## Introduction

Coronaviruses (CoVs) are members of the subfamily Orthocoronavirinae in the family Coronaviridae (*1*), and consist of four genera, *i.e*., alpha-, beta-, gamma- and delta-coronaviruses, as classified by the International Committee on Taxonomy of Viruses (ICTV) (*2-5*). All CoVs characterized so far are known to only infect vertebrates, and lead to various zoonotic diseases. Among them, three beta-CoVs are responsible for three major outbreaks of this century (*6-10*), *i.e*., the severe acute respiratory syndrome (SARS), the Middle East respiratory syndrome (MERS), and the ongoing global COVID-19 pandemic. Several alpha-CoVs and gamma-CoVs can also greatly affect livestock (*11-13*). For example, in 2017, the swine acute diarrhea syndrome coronavirus, an alpha-CoV, led to the death of more than 24,000 piglets in Guangdong, China (*11*).

Characteristic features of CoVs include a high genetic diversity (*7, 14*), frequent mutation (*15*), recombination (*16, 17*), and most importantly, frequent cross-host transmissions (*11, 18-20*) which is facilitated by the spike protein (*21-23*). These features pose serious challenges for the eradication and mitigation campaigns in case of epidemics, and make even the taxonomic classification of these viruses a challenging task. The traditional classification of CoVs (*2-5*) relies on phenotypic information such as the host range, and takes into account of the phylogeny of the viruses. Most of these phylogenetic analyses performed on CoVs rely on one of these two approaches (*3*) : (i) analyses based on the whole genome sequences; (ii) analyses based on individual conserved proteins, most prominently ORF1a/1b, spike, envelope, nucleocapsid, and membrane proteins. These analyses are useful for closely related species/lineages, but become problematic for more distant relationships (*24*). For example, the node robustness of phylogenetic trees built on the basis of whole genomes of distantly related species is often unreliable as can be judged by their bootstrap values (*25*). As a result, no comprehensive classification of coronaviruses is currently available.

In this study, we adopted a different approach based on the identification and use of a set of five concatenated CoVs protein clusters to create an Orthocoronavirinae species tree, by using the Markov Cluster Algorithm (MCL, https://www.micans.org/mcl/) (*26*). This phylogenetic species tree is robust, and is consistent with the current ICTV classification, and thus should be useful for the classification of future new CoV isolates. Furthermore, through reconciliation with the corresponding gene trees (*27-29*), this species tree allowed us to infer the apparent “horizontal gene transfer” (HGT) events for each of the five protein clusters, which we surmised can serve as a marker for cross-host transmissions. The specific cases of SARS-CoV, MERS-CoV, and SARS-CoV-2 are discussed.

## Results and Discussion

We searched through 3,880 genomes of coronaviruses from three databases NCBI, NGDC, and GISAID (https://www.ncbi.nlm.nih.gov/; https://bigd.big.ac.cn/; https://www.gisaid.org), and compiled 396 complete genomes with sequence identity lower than 95% (Supplementary Table S1). The predicted protein sequences were fed to the subsequent MCL analysis using graph clustering (*26*). In total, 116 protein clusters were identified, among which five clusters, i.e., ORF1b, ORF1a, spike, membrane, and nucleocapsid proteins were found to be the most prevalent (Table 1, Supplementary Fig. S1). These clusters were then chosen for alignment to construct a bootstrapped phylogenetic tree using Randomized Axelerated Maximum Likelihood (RAxML) (*30*) (Fig. 1). In this step, we added other 26 genome sequences isolated from different hosts, albeit with identity greater than 95% to the 396 sequences mentioned above (Supplementary Table S2). The bovine torovirus (NC_007447) from Tobaniviridae (*31, 32*) was selected as the outgroup. The virus genera (according to ICTV classification), the host families and host receptors (if available), and the size of genomes are also included in Fig. 1, in order to provide a comprehensive view of these metadata (*3, 4*).

**Table 1.** The five protein clusters as the single-copy markers of the subfamily Orthocoronavirinae.

| **Cluster Id** | **Number of**<br>**Single Copy Gene** | **Average**<br>**Length (aa)** | **Cluster Protein**<br>**Annotation** |
| --- | --- | --- | --- |
| Cluster1 | 397 | 2612 | ORF1b Protein |
| Cluster2 | 396 | 4103 | ORF1a Protein |
| Cluster3 | 396 | 1285 | Spike Protein |
| Cluster4 | 396 | 229 | Membrane Protein |
| Cluster5 | 396 | 405 | Nucleocapsid Protein |

**Figure 1.**
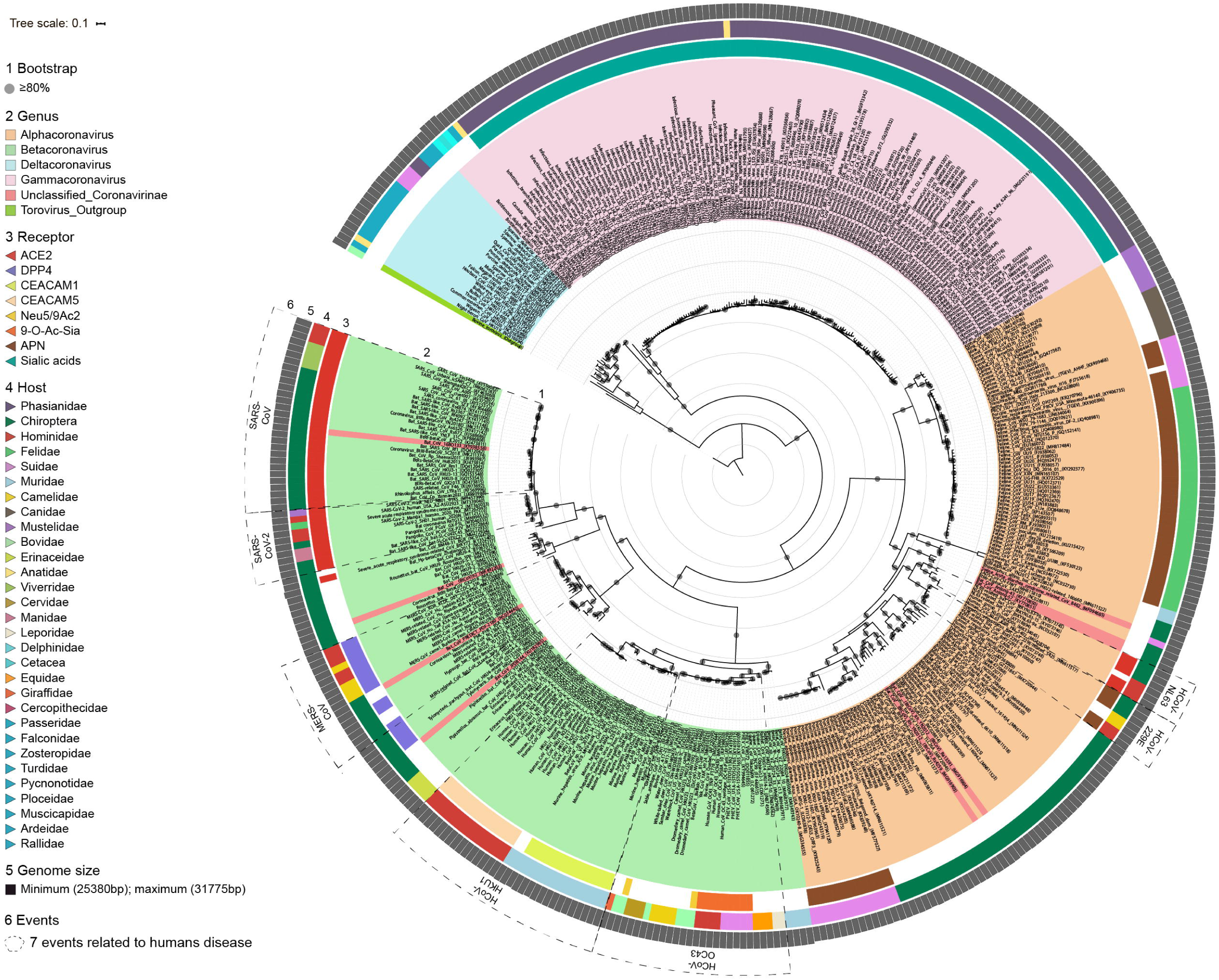
Rooted phylogeny of 422 Orthocoronavirinae viruses with the bovine torovirus from Tobaniviridae as the outgroup, inferred from five concatenated protein clusters using RAxML v7.2.8. Bootstrap support values ≥ 80% are shown as dots on interior nodes. Ring colors indicate the virus genus classification (indicated as squares in the legend) and their host phylogeny (indicated as triangles in the legend). Virus genome sizes are shown in the peripheral circle. Coronavirus information is provided in Supplementary Tables S1/S2. The visualization of the phylogenetic tree was performed by using iToL (https://itol.embl.de/) (*50*).

As can be seen in Fig. 1, this analysis yields genera and species that are largely congruous with the current ICTV classification (*5*), but the interpretation of the results is far more straightforward and requires just genome sequences. The resulting phylogenetic tree has a high degree of robustness, compared to those obtained using whole genome sequences, or the ORF1a/1b proteins or domains (see Methods). For example, nodes of branches of the most ancestral lineages of the tree were supported by bootstrap values ≥80%. Moreover, this phylogenetic tree can clearly distinguish different species of the viruses for the subfamily Orthocoronavirinae, while the phylogeny based on the ORF1a/b domains alone, which is better suited for classification at the order level (*33*), cannot do it.

Notably, this circular map format clearly shows the viruses along the same or closely related evolutionary branches can infect different hosts. For instance, in the specific case of SARS-CoV, 29 viruses clustered into a monophyletic clade, which included 22 strains isolated from Chiroptera (bats), four from Viverridae (palm civets), and three from Hominidae (humans). Similarly, several MERS-CoVs that form a monophyletic clade in the inferred tree were isolated from Chiroptera, Camelidae (dromedary camels), and Hominidae. In the section containing the coronaviruses Human_CoV_OC43 which are responsible for a common human flu, several hosts appear in the map (*34, 35*), reflecting the considerable promiscuity of their hosts including Bovidae (cattle), Leporidae (rabbits), Camelidae, Suidae (swine), and Equidae (horses).

We next determined the apparent HGT events among the viruses, which we assumed would indicate their cross-host transmissions. To this end, the reconciliations between the concatenated phylogeny and the related gene trees for the five protein clusters was performed, using RANGER-DTL version 2.0 (*36*). A visualized summary is given in Fig. 2 (*30*). We identified three types of transfer: (i) clade to clade, (ii) clade to strain, (iii) strain to strain. Within the same host, HGT occurred most frequently for Phasianidae, followed by Felidae and Hominidae (Fig. 2B). Transfer events across different hosts for clusters 3, 4, 5 (spike, membrane, and nucleocapsid proteins, respectively) were more frequent than for the other two clusters. Among different hosts, bats were the most frequent donor or receiver, followed by Suidae, Camelidae, Hominidae, and Phasianidae (Fig. 2C). This immediately suggests an urgent need for erecting ecological barriers between livestock and wild animals, and between humans and wild animals, as well as for worldwide monitoring of coronaviruses.

**Figure 2.**
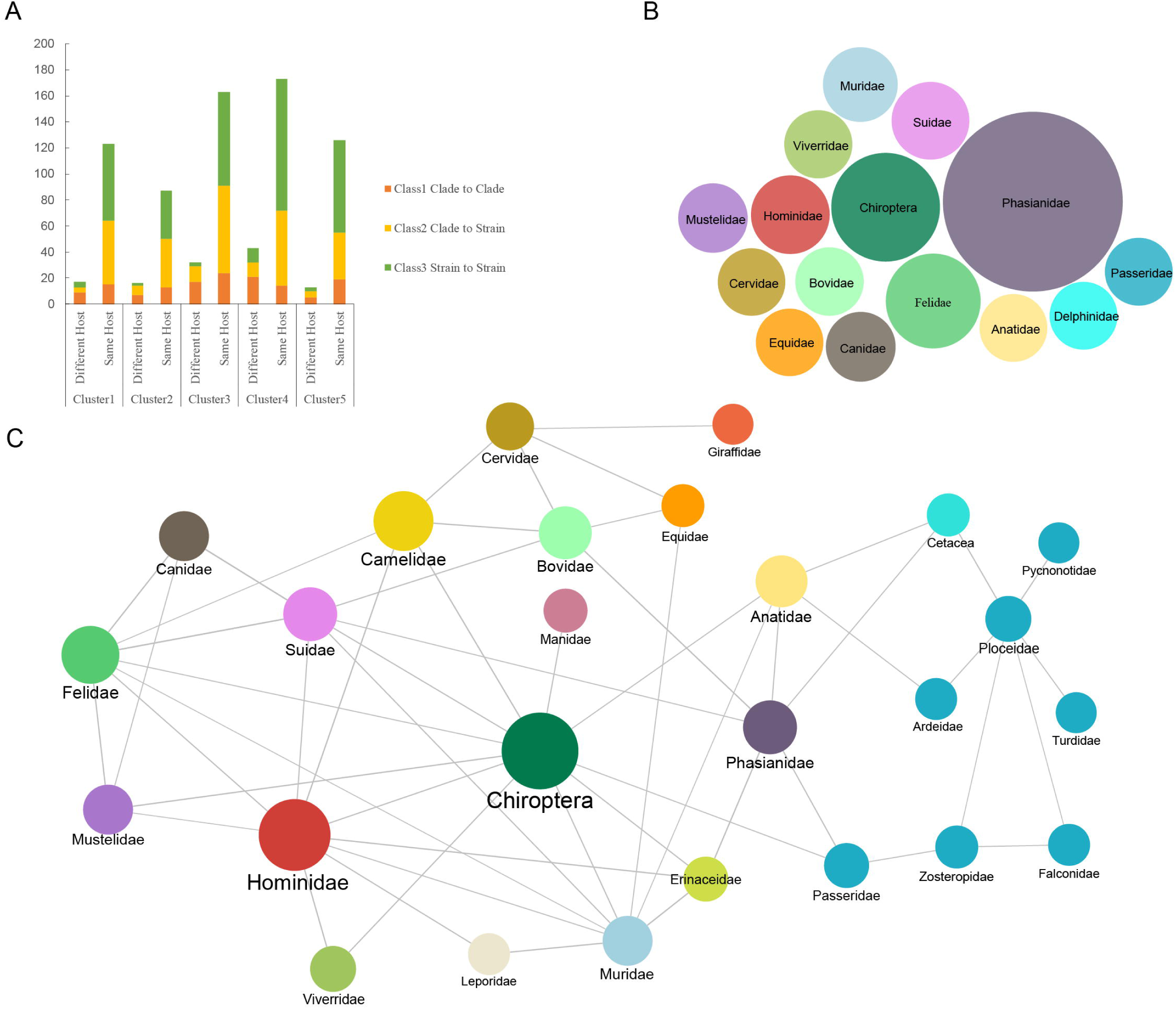
Statistics of the inferred horizontal gene transfer (HGT) events and host jumps of coronaviruses. A. Counts of transfer events that are within the same host family and among different host families. The colors within a bar stand for three different type of transfer events (i.e., from clade to clade, from clade to strain, from strain to strain. B. Counts of transfer events within the same host family. Each circle indicates a host family as shown in the label and its size is proportional to the number of transfer events within the same host family. C. Inferred host jump network of coronaviruses based on the transfer events. Each line indicates the predicted host jump between two host families and its thickness is proportional to the number of host jump events. Each circle indicates host family as shown in the label and its size is proportional to the number of host jump between different host families. (Figs. 2B and 2C is generated using Gephi 0.9.2, https://gephi.org) (*30*).

We then examined in more details the inferred HGT events for SARS-CoV, MERS-CoV and SARS-CoV-2. For this analysis, an expanded dataset of 269 complete genomes of the beta-CoVs with sequence identity lower than 99.8% was used, except that only four SARS-CoV-2 genomes of human origin were included (*37*), to reduce the redundancy (Supplementary Table S3). The Human_CoV_229E (NC_002645) from alpha-CoVs (*3*) was selected as the outgroup. For SARS-CoV (Fig. 3A), there were apparent transfer links among Chiroptera, Viverridae and Hominidae. This result is consistent with the current knowledge concerning the evolution of SARS-CoVs, with the palm civet as the likely intermediate host (*16, 38, 39*). As there is no evidence for direct transmission of this virus between Chiroptera and Hominidae, the gene transfer between Hominidae and Chiroptera as depicted in Fig. 3A might be an indirect result of separated transfer events between Chiroptera and Viverridae, and between Viverridae and Hominidae. The pattern found for MERS-CoV (Fig. 3B) is again consistent with accumulated evidence indicating the dromedary camel as a probable vector for human infections (*8, 40, 41*). For SARS-CoV-2 (Fig. 3C), transfer links were found between Chiroptera and Hominidae, and between Chiroptera and Manidae (pangolins). Interestingly, no gene transfer between Hominidae and Manidae can be inferred from our analysis. Pangolins were suggested to be involved in the species jump of SARS-CoV-2 to humans (*37, 42*). However, more recent studies have disputed this notion (*37, 43, 44*). Finally, while the links among Hominidae, Mustelidae (minks), and Felidae (tigers) were deduced, it remains to be seen how SARS-CoV-2 viruses were transmitted among them. Taken together, we argue that these inferred apparent HGT events can serve as a possible marker for cross-host transmissions of coronaviruses.

**Figure 3.**
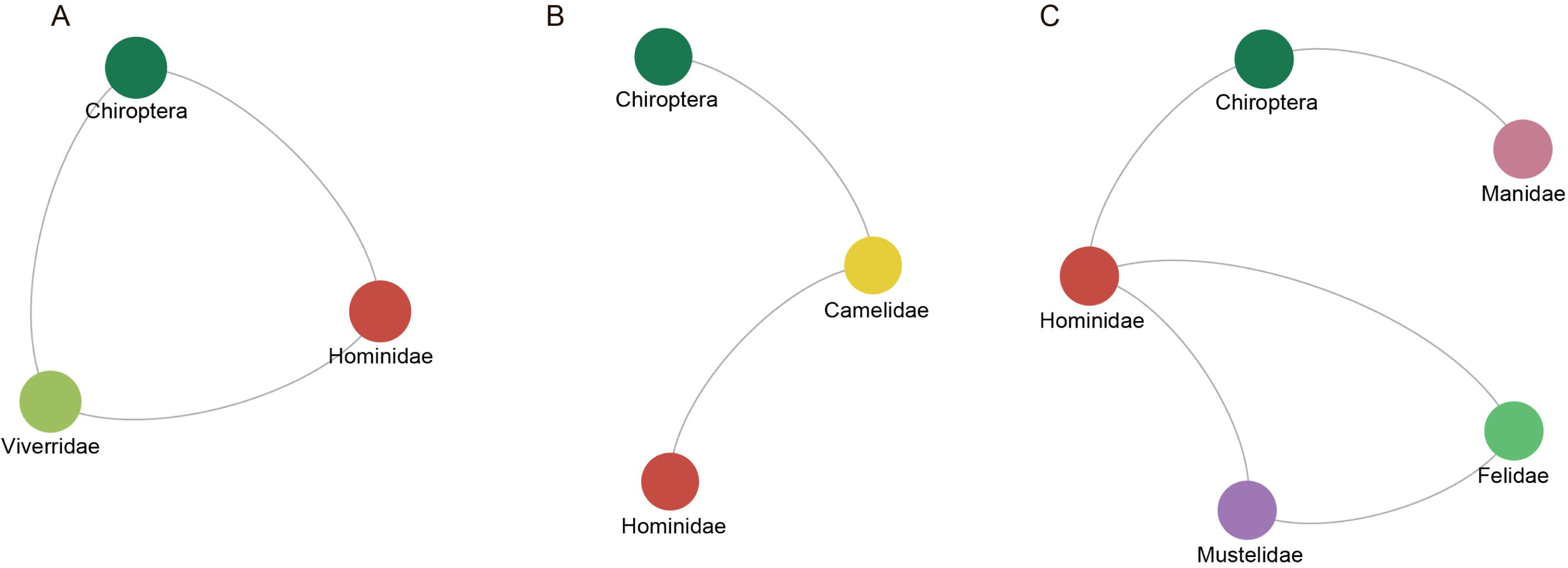
Cross-host transmissions inferred from horizontal gene transfer (HGT) events for SARS-CoV (A), MERS-CoV (B), and SARS-CoV-2 (C). The circles correspond to the hosts, and the lines correspond to the apparent HGT events among the hosts.

In summary, based on the analysis of concatenated protein clusters of the viral genome sequences, we present a framework to produce robust phylogeny and classification for CoVs, and to infer the apparent HGT events and cross host transmissions for the Orthocoronavirinae subfamily. This should help to understand how CoVs are transmitted to humans, and aid in devising strategies to combat the severe global health threat posed by these viruses.

## Methods

### Genome sequence collection and marker selection

Genome datasets were compiled with CD-HIT version 4.7 (*45*). Open reading frames (ORFs) in the datasets were predicted by using GeneMarkS version 4.32 (*46*), and then annotated using BLAST against the NR database (https://ftp.ncbi.nlm.nih.gov/blast/db/FASTA/) with e-value ≤ 10^−5^.

Marker selections were performed with the Markov Cluster algorithm of OrthoMCL, with the parameter of ‘-I 1.5’ (*26*). The 20 protein clusters with abundancy in a genus higher than 30% are listed in Supplementary Fig. S1.

### Phylogenetic inference and apparent HGT inference

For the phylogenetic analyses, the multiple sequence alignments (MSAs) of the datasets (the 422 genome sequences used for the analysis of the subfamily Orthocoronavirinae, and the 269 genome sequences used for the analysis of beta-CoVs) were analyzed by MAFFT v7.407 (*47*) based on the five concatenated protein clusters or each individual cluster. Subsequently, maximum likelihood phylogenies were estimated using RAxML version 7.2.8 (*48*), utilizing the PROTGAMMALG model with 100 bootstrap replicates. The apparent HGT events were inferred based on reconciliation of the species tree and the corresponding gene tree (*27*), by using RANGER-DTL version 2.0 with default parameters (https://compbio.engr.uconn.edu/software/RANGER-DTL/) (*36, 49*). The apparent HGT events were then presented by using Gephi 0.9.2.

For comparison of robustness, two other phylogenetic trees were also built, based on the full genome, and the five concatenated domains of ORF1a/b that are used in the ICTV classification approach by the Coronaviridae Study Group (*e.g*., 3CLpro, NiRAN, RdRp, ZBD, and HEL1) (*3, 33*). The phylogenetic tree based on the full genome was constructed using RAxML with the GTRGAMMA model and 1,000 bootstrap replicates. The phylogenetic tree based on the five concatenated domains of ORF1a/1b was built using the same method as that based on the five concatenated protein clusters described above.

