## Supplementary Table S1, Supplementary Table S2, Supplementary Table S3, Supplementary Figure S1 for "A Comprehensive Classification of Coronaviruses and Inferred Cross-Host Transmissions"

**Supplementary Materials**

**Supplementary Table S1** The 396 coronavirus genomes used in the identification of protein clusters

| **Accession ID** | **Strain** | **Genus** | **Host** | **Length (bp)** |
| --- | --- | --- | --- | --- |
| AB551247 | Murine_hepatitis_virus_MHV-MI | Betacoronavirus | Muridae | 31428 |
| AC_000192 | Murine_hepatitis_virus_JHM | Betacoronavirus | Muridae | 31526 |
| AF201929 | Murine_hepatitis_virus_2 | Betacoronavirus | Muridae | 31276 |
| AF208066 | Murine_hepatitis_virus_Penn_97-1 | Betacoronavirus | Muridae | 31112 |
| AF208067 | Murine_hepatitis_virus_ML-10 | Betacoronavirus | Muridae | 31233 |
| AF220295 | Bovine_CoV_Quebec | Betacoronavirus | Bovidae | 31100 |
| AY319651 | Avian_infectious_bronchitis_virus | Gammacoronavirus | Phasianidae | 27733 |
| AY463060 | SARS_CoV_ShanghaiQXC2 | Betacoronavirus | Hominidae | 29013 |
| AY514485 | Infectious_bronchitis_virus_serotype_California_99 | Gammacoronavirus | Phasianidae | 27693 |
| AY518894 | Human_group_1_CoV | Alphacoronavirus | Hominidae | 27555 |
| AY559083 | SARS_CoV_Sin3408 | Betacoronavirus | Hominidae | 29767 |
| AY585229 | Human_CoV_OC43_serotype_OC43-Paris | Betacoronavirus | Hominidae | 30744 |
| AY641576 | Avian_infectious_bronchitis_virus | Gammacoronavirus | Phasianidae | 27434 |
| AY884001 | Human_CoV_HKU1_genotype_B | Betacoronavirus | Hominidae | 29815 |
| DQ010921 | Feline_CoV_FIPV_79-1146 | Alphacoronavirus | Felidae | 29147 |
| DQ011855 | PHEV_VW572 | Betacoronavirus | Suidae | 30480 |
| DQ022305 | Bat_SARS_CoV_HKU3-1 | Betacoronavirus | Chiroptera | 29728 |
| DQ288927 | Avian_infectious_bronchitis_virus | Gammacoronavirus | Phasianidae | 27534 |
| DQ339101 | Human_CoV_HKU1_N5P8_genotype_A_B | Betacoronavirus | Hominidae | 29755 |
| DQ412042 | Bat_SARS_CoV_Rf1 | Betacoronavirus | Chiroptera | 29709 |
| DQ412043 | Bat_SARS_CoV_Rm1 | Betacoronavirus | Chiroptera | 29749 |
| DQ415896 | Human_CoV_HKU1_N19_genotype_A | Betacoronavirus | Hominidae | 29860 |
| DQ415897 | Human_CoV_HKU1_N20_genotype_C | Betacoronavirus | Hominidae | 29845 |
| DQ415899 | Human_CoV_HKU1_N22_genotype_C | Betacoronavirus | Hominidae | 29905 |
| DQ415900 | Human_CoV_HKU1_N23_genotype_A | Betacoronavirus | Hominidae | 29812 |
| DQ415901 | Human_CoV_HKU1_N24_genotype_A | Betacoronavirus | Hominidae | 30097 |
| DQ415904 | Human_CoV_HKU1_N6_genotype_A | Betacoronavirus | Hominidae | 29295 |
| DQ415908 | Human_CoV_HKU1_N11_genotype_A | Betacoronavirus | Hominidae | 29947 |
| DQ415909 | Human_CoV_HKU1_N13_genotype_A | Betacoronavirus | Hominidae | 29767 |
| DQ415913 | Human_CoV_HKU1_N17_genotype_C | Betacoronavirus | Hominidae | 29785 |
| DQ811786 | TGEV_Miller_M60 | Alphacoronavirus | Suidae | 27904 |
| DQ811787 | PRCV_ISU-1 | Alphacoronavirus | Suidae | 27550 |
| DQ848678 | Feline_CoV_FCoV_C1Je | Alphacoronavirus | Felidae | 29277 |
| EF065508 | Bat_CoV_HKU4-4 | Betacoronavirus | Chiroptera | 30316 |
| EF065514 | Bat_CoV_HKU9-2 | Betacoronavirus | Chiroptera | 29107 |
| EF065516 | Bat_CoV_HKU9-4 | Betacoronavirus | Chiroptera | 29155 |
| EF424623 | Giraffe_CoV_US_OH3 | Betacoronavirus | Giraffidae | 31002 |
| EF446615 | Equine_CoV_NC99 | Betacoronavirus | Equidae | 30992 |
| EU022525 | Turkey_CoV(TCoV)-540 | Gammacoronavirus | Phasianidae | 27771 |
| EU022526 | Turkey_CoV(TCoV)-ATCC | Gammacoronavirus | Phasianidae | 27854 |
| EU186072 | Feline_CoV | Alphacoronavirus | Felidae | 29256 |
| EU418975 | Infectious_bronchitis_virus_ArkDPI101 | Gammacoronavirus | Phasianidae | 27636 |
| EU420137 | Bat_CoV_1B_AFCD307 | Alphacoronavirus | Chiroptera | 28476 |
| EU526388 | Infectious_bronchitis_virus_A2 | Gammacoronavirus | Phasianidae | 27715 |
| EU637854 | Infectious_bronchitis_virus_LSD_05I | Gammacoronavirus | Phasianidae | 27638 |
| FJ588686 | Bat_SARS_CoV_Rs672 | Betacoronavirus | Chiroptera | 29059 |
| FJ647222 | Murine_CoV_SA59_RJHM | Betacoronavirus | Muridae | 31283 |
| FJ647223 | Murine_CoV_MHV-1 | Betacoronavirus | Muridae | 31386 |
| FJ647224 | Murine_CoV_MHV-3 | Betacoronavirus | Muridae | 31448 |
| FJ755618 | Transmissible_gastroenteritis_virus_H16 | Alphacoronavirus | Suidae | 28569 |
| FJ904714 | Infectious_bronchitis_virus_Cal | Gammacoronavirus | Phasianidae | 27664 |
| FJ904715 | Infectious_bronchitis_virus_Cal557 | Gammacoronavirus | Phasianidae | 27615 |
| FJ904722 | Infectious_bronchitis_virus_Mass41 | Gammacoronavirus | Phasianidae | 27614 |
| FJ938051 | Feline_CoV_RM | Alphacoronavirus | Felidae | 29232 |
| FJ938052 | Feline_CoV_UU11 | Alphacoronavirus | Felidae | 29306 |
| FJ938053 | Feline_CoV_UU7 | Alphacoronavirus | Felidae | 29277 |
| FJ938056 | Feline_CoV_UU5 | Alphacoronavirus | Felidae | 29253 |
| FJ938057 | Feline_CoV_UU15 | Alphacoronavirus | Felidae | 29275 |
| FJ938058 | Feline_CoV_UU16 | Alphacoronavirus | Felidae | 28479 |
| FJ938061 | Feline_CoV_UU3 | Alphacoronavirus | Felidae | 29130 |
| FJ938062 | Feline_CoV_UU9 | Alphacoronavirus | Felidae | 29266 |
| FN430414 | Infectious_bronchitis_virus_ITA_90254 | Gammacoronavirus | Phasianidae | 27643 |
| FN430415 | Infectious_bronchitis_virus_NGA_A116E7 | Gammacoronavirus | Phasianidae | 27593 |
| GQ152141 | Feline_CoV_FCoV_NTU156_P | Alphacoronavirus | Felidae | 28897 |
| GQ153543 | Bat_SARS_CoV_HKU3-8 | Betacoronavirus | Chiroptera | 29681 |
| GQ153548 | Bat_SARS_CoV_HKU3-13 | Betacoronavirus | Chiroptera | 29677 |
| GQ427175 | Turkey_CoV_TCoV_IN-517_94 | Gammacoronavirus | Phasianidae | 27690 |
| GQ427176 | Turkey_CoV_TCoV_TX-1038_98 | Gammacoronavirus | Phasianidae | 27804 |
| GQ477367 | Canine_CoV_CCoV_NTU336_F | Alphacoronavirus | Canidae | 29363 |
| GQ504720 | Infectious_bronchitis_virus_Arkansas_DPI | Gammacoronavirus | Phasianidae | 27652 |
| GQ504722 | Infectious_bronchitis_virus | Gammacoronavirus | Phasianidae | 27648 |
| GQ504723 | Infectious_bronchitis_virus | Gammacoronavirus | Phasianidae | 27620 |
| GQ504725 | Infectious_bronchitis_virus_Mass41_Vaccine | Gammacoronavirus | Phasianidae | 27451 |
| GU393332 | Infectious_bronchitis_virus_serotype_Delaware_072 | Gammacoronavirus | Phasianidae | 27591 |
| GU393333 | Infectious_bronchitis_virus_serotype_FL18288 | Gammacoronavirus | Phasianidae | 27616 |
| GU393334 | Infectious_bronchitis_virus_serotype_Gray | Gammacoronavirus | Phasianidae | 27622 |
| GU393337 | Infectious_bronchitis_virus_serotype_Iowa_97 | Gammacoronavirus | Phasianidae | 27663 |
| GU553361 | Feline_CoV_UU22 | Alphacoronavirus | Felidae | 29264 |
| HM211098 | Bat_CoV_HKU9-5-1 | Betacoronavirus | Chiroptera | 29136 |
| HM211101 | Bat_CoV_HKU9-10-2 | Betacoronavirus | Chiroptera | 29122 |
| HM245924 | Infectious_bronchitis_virus | Gammacoronavirus | Phasianidae | 27709 |
| HM245926 | Mink_CoV_WD1133 | Alphacoronavirus | Mustelidae | 28915 |
| HQ012367 | Feline_CoV_UU17 | Alphacoronavirus | Felidae | 29202 |
| HQ012369 | Feline_CoV_UU21 | Alphacoronavirus | Felidae | 28429 |
| HQ012370 | Feline_CoV_UU24 | Alphacoronavirus | Felidae | 29266 |
| HQ012371 | Feline_CoV_UU31 | Alphacoronavirus | Felidae | 29274 |
| HQ392470 | Feline_CoV_UU19 | Alphacoronavirus | Felidae | 29255 |
| HQ392471 | Feline_CoV_UU20 | Alphacoronavirus | Felidae | 29252 |
| HQ848267 | Infectious_bronchitis_virus_GX-YL5 | Gammacoronavirus | Phasianidae | 27720 |
| JF274479 | Infectious_bronchitis_virus_LHLJ_07VII | Gammacoronavirus | Phasianidae | 27678 |
| JF705860 | Duck_CoV_DK_HN_ZZ2004 | Gammacoronavirus | Anatidae | 27673 |
| JF792616 | Rat_CoV_681 | Betacoronavirus | Muridae | 31286 |
| JF893452 | Infectious_bronchitis_virus_YN | Gammacoronavirus | Phasianidae | 27681 |
| JN183882 | Feline_CoV_UU47 | Alphacoronavirus | Felidae | 29243 |
| JN183883 | Feline_CoV_UU54 | Alphacoronavirus | Felidae | 29222 |
| JN634064 | Feline_CoV_WSU_79-1683 | Alphacoronavirus | Felidae | 28984 |
| JN856008 | Canine_CoV_A76 | Alphacoronavirus | Canidae | 29271 |
| JN874560 | Rabbit_CoV_HKU14_HKU14-3 | Betacoronavirus | Leporidae | 31116 |
| JN874562 | Rabbit_CoV_HKU14_HKU14-10 | Betacoronavirus | Leporidae | 30904 |
| JQ088078 | Infectious_bronchitis_virus_CK_SWE_0658946_10 | Gammacoronavirus | Phasianidae | 27664 |
| JQ173883 | Murine_hepatitis_virus_S_3239-17 | Betacoronavirus | Muridae | 31161 |
| JQ404409 | Canine_CoV_1-71 | Alphacoronavirus | Canidae | 29462 |
| JQ404410 | Canine_CoV_TN-449 | Alphacoronavirus | Canidae | 29364 |
| JQ408980 | Feline_CoV_DF-2_R3i | Alphacoronavirus | Felidae | 29363 |
| JQ408981 | Feline_infectious_peritonitis_virus_DF-2 | Alphacoronavirus | Felidae | 29025 |
| JQ977697 | Infectious_bronchitis_virus_SNU8067 | Gammacoronavirus | Phasianidae | 27708 |
| JQ989269 | Hipposideros_bat_CoV_HKU10_LSH5A | Alphacoronavirus | Chiroptera | 28492 |
| JX169866 | Murine_CoV_JHM-WU-Dns2 | Betacoronavirus | Muridae | 30729 |
| JX169867 | Murine_CoV_JHM.WU | Betacoronavirus | Muridae | 31051 |
| JX195178 | Infectious_bronchitis_virus_LDL_97I_substrain_P115 | Gammacoronavirus | Phasianidae | 27693 |
| JX993987 | Bat_CoV_Rp_Shaanxi2011 | Betacoronavirus | Chiroptera | 29484 |
| JX993988 | Bat_CoV_Cp_Yunnan2011 | Betacoronavirus | Chiroptera | 29452 |
| KC008600 | Infectious_bronchitis_virus | Gammacoronavirus | Phasianidae | 27701 |
| KC013541 | Infectious_bronchitis_virus_LGD_120723 | Gammacoronavirus | Phasianidae | 27718 |
| KC506155 | Infectious_bronchitis_virus_LJL_111054 | Gammacoronavirus | Phasianidae | 27648 |
| KC545386 | Erinaceus_CoV_2012-216_GER | Betacoronavirus | Erinaceidae | 30175 |
| KC869678 | Coronavirus_Neoromicia_PML-PHE1_RSA | Betacoronavirus | Chiroptera | 30111 |
| KC881006 | Bat_SARS-like_CoV_Rs3367 | Betacoronavirus | Chiroptera | 29792 |
| KF411040 | Infectious_bronchitis_virus_LLN_111169 | Gammacoronavirus | Phasianidae | 27663 |
| KF460437 | Infectious_bronchitis_virus_VicS-v | Gammacoronavirus | Phasianidae | 27610 |
| KF530123 | Feline_CoV_Felis_catus_NLD_UU88 | Alphacoronavirus | Felidae | 29123 |
| KF569996 | Rhinolophus_affinis_CoV_LYRa11 | Betacoronavirus | Chiroptera | 29805 |
| KF686343 | HKU1_human_USA_HKU1-13 | Betacoronavirus | Hominidae | 29983 |
| KF686344 | HKU1_human_USA_HKU1-15 | Betacoronavirus | Hominidae | 29695 |
| KF793826 | Bottlenose_dolphin_CoV_HKU22_CF090331 | Gammacoronavirus | Delphinidae | 31775 |
| KJ425503 | Infectious_bronchitis_virus_LHLJ_090908 | Gammacoronavirus | Phasianidae | 27609 |
| KJ435283 | Infectious_bronchitis_virus_LSD_111219 | Gammacoronavirus | Phasianidae | 27666 |
| KJ473797 | BtMf-AlphaCoV_GD2012 | Alphacoronavirus | Chiroptera | 28758 |
| KJ473798 | BtMf-AlphaCoV_HuB2013 | Alphacoronavirus | Chiroptera | 28755 |
| KJ473799 | BtMf-AlphaCoV_FJ2012 | Alphacoronavirus | Chiroptera | 28765 |
| KJ473810 | BtMs-AlphaCoV_GS2013 | Alphacoronavirus | Chiroptera | 27576 |
| KJ473811 | BtRf-BetaCoV_JL2012 | Betacoronavirus | Chiroptera | 29037 |
| KJ473814 | BtRs-BetaCoV_HuB2013 | Betacoronavirus | Chiroptera | 29658 |
| KJ473815 | BtRs-BetaCoV_GX2013 | Betacoronavirus | Chiroptera | 29161 |
| KJ473821 | BtVs-BetaCoV_SC2013 | Betacoronavirus | Chiroptera | 30423 |
| KM213963 | Infectious_bronchitis_virus_XDC_2 | Gammacoronavirus | Phasianidae | 27714 |
| KM349743 | BetaCoV_HKU24_HKU24-R05009I | Betacoronavirus | Muridae | 31249 |
| KM454473 | Duck_CoV_DK_GD_27 | Gammacoronavirus | Anatidae | 27754 |
| KP118887 | Infectious_bronchitis_virus_LHB_140532 | Gammacoronavirus | Phasianidae | 27621 |
| KP118892 | Infectious_bronchitis_virus_LLN_130101 | Gammacoronavirus | Phasianidae | 27594 |
| KP118894 | Infectious_bronchitis_virus_LGD_090907 | Gammacoronavirus | Phasianidae | 27685 |
| KP143507 | Feline_CoV_27C | Alphacoronavirus | Felidae | 28984 |
| KP198610 | Human_CoV_OC43_2058A_10 | Betacoronavirus | Hominidae | 30752 |
| KP662631 | Infectious_bronchitis_virus_3665_11 | Gammacoronavirus | Phasianidae | 27388 |
| KP790143 | Infectious_bronchitis_virus_LDL_140520 | Gammacoronavirus | Phasianidae | 27712 |
| KP849472 | AlphaCoV_1_23_03 | Alphacoronavirus | Canidae | 30004 |
| KP868572 | Infectious_bronchitis_virus_LHLJ_111043 | Gammacoronavirus | Phasianidae | 27452 |
| KP886808 | Bat_SARS-like_CoV_YNLF_31C | Betacoronavirus | Chiroptera | 29723 |
| KP887098 | Murine_CoV_AM2 | Betacoronavirus | Muridae | 31409 |
| KP981644 | Canine_CoV_CB_05 | Alphacoronavirus | Canidae | 29278 |
| KR095279 | Porcine_epidemic_diarrhea_virus_HNQX-3_14 | Alphacoronavirus | Suidae | 27997 |
| KR231009 | Infectious_bronchitis_virus_B1648 | Gammacoronavirus | Phasianidae | 27654 |
| KR270796 | Porcine_respiratory_CoV_OH7269 | Alphacoronavirus | Suidae | 27765 |
| KR608272 | Infectious_bronchitis_virus_LDT3-A | Gammacoronavirus | Phasianidae | 27681 |
| KR822424 | European_turkey_CoV_080385d | Gammacoronavirus | Phasianidae | 27739 |
| KR902510 | Infectious_bronchitis_virus_Ind-TN92-03 | Gammacoronavirus | Phasianidae | 27464 |
| KT368891 | Camel_CoV_HKU23_camel | Betacoronavirus | Camelidae | 31041 |
| KT368894 | Camel_alphaCoV_camel | Alphacoronavirus | Camelidae | 27397 |
| KT779555 | Human_CoV_HKU1_BJ01-p3 | Betacoronavirus | Hominidae | 29887 |
| KT852992 | Infectious_bronchitis_virus_tl_LDT3_03 | Gammacoronavirus | Phasianidae | 27699 |
| KT886454 | Infectious_bronchitis_virus_(gammaCoV)_74 | Gammacoronavirus | Phasianidae | 27418 |
| KT941120 | Porcine_epidemic_diarrhea_virus_HUA-14PED96 | Alphacoronavirus | Suidae | 27966 |
| KU182965 | Bat_CoV_JPDB144 | Unclassified_Coronavirinae | Chiroptera | 30321 |
| KU215419 | Feline_CoV_inoculum | Alphacoronavirus | Felidae | 29262 |
| KU215427 | Feline_CoV_Cat3_day28_deletion | Alphacoronavirus | Felidae | 29238 |
| KU317090 | Infectious_bronchitis_virus_SAIBK2 | Gammacoronavirus | Phasianidae | 27669 |
| KU356856 | Infectious_bronchitis_virus_SCYB_140913 | Gammacoronavirus | Phasianidae | 27647 |
| KU361188 | Infectious_bronchitis_virus_2014_QL1403 | Gammacoronavirus | Phasianidae | 27691 |
| KU710265 | MERS_CoV_Hu_Taif_KSA-7032_2014_S530del | Betacoronavirus | Hominidae | 29566 |
| KU851859 | MERS_CoV_Hu_Jeddah-KSA-3RS2702 | Betacoronavirus | Hominidae | 30076 |
| KU900739 | Infectious_bronchitis_virus_QIA-03342 | Gammacoronavirus | Phasianidae | 27684 |
| KU973692 | SARS-related_CoV_F46 | Betacoronavirus | Chiroptera | 29722 |
| KX077987 | Infectious_bronchitis_virus_LDL_150434-III | Gammacoronavirus | Phasianidae | 27666 |
| KX185057 | Infectious_bronchitis_virus_LHLJ_95I | Gammacoronavirus | Phasianidae | 27802 |
| KX185059 | Infectious_bronchitis_virus_LH1 | Gammacoronavirus | Phasianidae | 27681 |
| KX219791 | Infectious_bronchitis_virus_LHLJ_07I | Gammacoronavirus | Phasianidae | 27648 |
| KX219797 | Infectious_bronchitis_virus_LXJ_02I | Gammacoronavirus | Phasianidae | 27684 |
| KX236001 | Infectious_bronchitis_virus_LSD_03I | Gammacoronavirus | Phasianidae | 27631 |
| KX236005 | Infectious_bronchitis_virus_LJL_08-1 | Gammacoronavirus | Phasianidae | 27657 |
| KX252778 | Infectious_bronchitis_virus_LJL_05I | Gammacoronavirus | Phasianidae | 27677 |
| KX252779 | Infectious_bronchitis_virus_LLN_06I | Gammacoronavirus | Phasianidae | 27684 |
| KX252782 | Infectious_bronchitis_virus_LLN_07I | Gammacoronavirus | Phasianidae | 27654 |
| KX252785 | Infectious_bronchitis_virus_LSD_100311 | Gammacoronavirus | Phasianidae | 27686 |
| KX252791 | Infectious_bronchitis_virus_LLN_98I | Gammacoronavirus | Phasianidae | 27683 |
| KX272465 | Infectious_bronchitis_virus_AR251-15 | Gammacoronavirus | Phasianidae | 27646 |
| KX302862 | Infectious_bronchitis_virus_LJS_101111 | Gammacoronavirus | Phasianidae | 27642 |
| KX302866 | Infectious_bronchitis_virus_LJL_04I | Gammacoronavirus | Phasianidae | 27714 |
| KX348114 | Infectious_bronchitis_virus_LDL_05III | Gammacoronavirus | Phasianidae | 27687 |
| KX364296 | Infectious_bronchitis_virus_LJL_08-9 | Gammacoronavirus | Phasianidae | 27655 |
| KX425847 | Infectious_bronchitis_virus_LJL_140734 | Gammacoronavirus | Phasianidae | 27709 |
| KX434790 | Infectious_bronchitis_virus_LJS_111210 | Gammacoronavirus | Phasianidae | 27618 |
| KX442564 | Hypsugo_bat_CoV_HKU25_YD131305 | Betacoronavirus | Chiroptera | 30498 |
| KX499468 | Transmissible_gastroenteritis_virus_(TGEV)_AHHF | Alphacoronavirus | Suidae | 28614 |
| KX512809 | Ferret_enteric_CoV_FECV1 | Alphacoronavirus | Mustelidae | 28151 |
| KX534205 | Porcine_epidemic_diarrhea_virus_JSLS-1 | Alphacoronavirus | Suidae | 27861 |
| KX722529 | Feline_CoV_UG-FH8 | Alphacoronavirus | Felidae | 29174 |
| KX722530 | Feline_CoV_Cat_1_Karlslunde | Alphacoronavirus | Felidae | 28595 |
| KX839246 | Porcine_epidemic_diarrhea_virus_85-7 | Alphacoronavirus | Suidae | 28015 |
| KX839248 | Porcine_epidemic_diarrhea_virus_85-7-mutant2 | Alphacoronavirus | Suidae | 27533 |
| KX900396 | Transmissible_gastroenteritis_virus_(TGEV) | Alphacoronavirus | Suidae | 28393 |
| KX981440 | Porcine_epidemic_diarrhea_virus | Alphacoronavirus | Suidae | 28124 |
| KY063617 | Canine_CoV_HLJ-072 | Alphacoronavirus | Canidae | 29264 |
| KY063618 | Canine_CoV_HLJ-073 | Alphacoronavirus | Canidae | 28969 |
| KY073745 | NL63-related_bat_CoV_BtKYNL63-9b | Alphacoronavirus | Chiroptera | 28679 |
| KY073746 | NL63-related_bat_CoV_BtKYNL63-15 | Alphacoronavirus | Chiroptera | 28442 |
| KY073747 | 229E-related_bat_CoV_BtKY229E-1 | Alphacoronavirus | Chiroptera | 27837 |
| KY073748 | 229E-related_bat_CoV_BtKY229E-8 | Alphacoronavirus | Chiroptera | 27636 |
| KY292377 | Feline_CoV_HLJ_DQ_2016_01 | Alphacoronavirus | Felidae | 29303 |
| KY352407 | Severe_acute_respiratory_syndrome-related_CoV_BtKY72 | Betacoronavirus | Chiroptera | 29274 |
| KY369909 | Human_CoV_229E_SC677 | Alphacoronavirus | Hominidae | 26592 |
| KY406735 | Porcine_respiratory_CoV_PRCV_USA_Minnesota-46140 | Alphacoronavirus | Suidae | 27779 |
| KY407556 | Infectious_bronchitis_virus_(gammaCoV)_I0114_14 | Gammacoronavirus | Phasianidae | 27681 |
| KY417145 | Bat_SARS-like_CoV_Rf4092 | Betacoronavirus | Chiroptera | 29710 |
| KY417148 | Bat_SARS-like_CoV_Rs4247 | Betacoronavirus | Chiroptera | 29743 |
| KY417150 | Bat_SARS-like_CoV_Rs4874 | Betacoronavirus | Chiroptera | 30311 |
| KY419109 | PHEV_CoV_USA-15TOSU1655 | Betacoronavirus | Suidae | 29906 |
| KY419110 | PHEV_CoV_USA-15TOSU1362 | Betacoronavirus | Suidae | 30515 |
| KY419112 | PHEV_CoV_USA-15TOSU1765 | Betacoronavirus | Suidae | 30531 |
| KY420075 | Porcine_epidemic_diarrhea_virus_PEDV-SX | Alphacoronavirus | Suidae | 27967 |
| KY566209 | Feline_CoV_HLJ_HRB_10 | Alphacoronavirus | Felidae | 29440 |
| KY620116 | Infectious_bronchitis_virus_(gammaCoV)_I1101_16 | Gammacoronavirus | Phasianidae | 27685 |
| KY626044 | Avian_CoV_BR-I | Gammacoronavirus | Phasianidae | 27618 |
| KY626045 | Avian_CoV_Ma5 | Gammacoronavirus | Phasianidae | 27652 |
| KY688119 | MERS-related_CoV_Hu_Aseer-KSA-Rs924 | Betacoronavirus | Hominidae | 30061 |
| KY770850 | Bat_CoV_Anlong-43 | Unclassified_Coronavirinae | Chiroptera | 26883 |
| KY770851 | Bat_CoV_Anlong-57 | Unclassified_Coronavirinae | Chiroptera | 28096 |
| KY799179 | Myotis_lucifugus_CoV | Alphacoronavirus | Chiroptera | 28173 |
| KY799582 | Infectious_bronchitis_virus_LSC_99I | Gammacoronavirus | Phasianidae | 27672 |
| KY805846 | Infectious_bronchitis_virus_IBV_Ck_EG_CU_4 | Gammacoronavirus | Phasianidae | 27663 |
| KY825243 | Porcine_epidemic_diarrhea_virus_KNU-141112-S_DEL5_ORF3 | Alphacoronavirus | Suidae | 27974 |
| KY933089 | Avian_CoV_1148-A | Gammacoronavirus | Phasianidae | 27573 |
| KY938558 | Bat_CoV_16BO133 | Unclassified_Coronavirinae | Chiroptera | 29075 |
| KY963963 | Porcine_epidemic_diarrhea_virus_KNU-1601 | Alphacoronavirus | Suidae | 28053 |
| KY994645 | PHEV_CoV_JL | Betacoronavirus | Suidae | 30684 |
| LC061272 | Equine_CoV_Tokachi09 | Betacoronavirus | Equidae | 30782 |
| LC061273 | Equine_CoV_Obihiro12-1 | Betacoronavirus | Equidae | 30916 |
| LC063838 | Porcine_epidemic_diarrhea_virus_MYG-1 | Alphacoronavirus | Suidae | 27481 |
| LC119077 | Ferret_CoV_FRCoV4370 | Alphacoronavirus | Mustelidae | 28550 |
| LC215871 | Ferret_CoV_ferret063 | Alphacoronavirus | Mustelidae | 28542 |
| LC364342 | Falcon_CoV_UAE-HKU27_988F | Deltacoronavirus | Falconidae | 26162 |
| LC364345 | Quail_CoV_UAE-HKU30_411F | Deltacoronavirus | Phasianidae | 25874 |
| LN610099 | Guinea_fowl_CoV_(GfCoV) | Gammacoronavirus | Phasianidae | 27471 |
| MF094685 | Swine_acute_diarrhea_syndrome_related_CoV_8462 | Unclassified_Coronavirinae | Chiroptera | 27200 |
| MF113046 | AlphaCoV_Minkina_1 | Alphacoronavirus | Mustelidae | 28924 |
| MF421319 | Infectious_bronchitis_virus_UY_09_CA_01 | Gammacoronavirus | Phasianidae | 27647 |
| MF421320 | Infectious_bronchitis_virus_UY_11_CA_18 | Gammacoronavirus | Phasianidae | 27634 |
| MF577027 | Porcine_epidemic_diarrhea_virus_(PEDV)_Belgorod_dom | Alphacoronavirus | Suidae | 28315 |
| MF618252 | Murine_hepatitis_virus_A59_WT-MHV_P250 | Betacoronavirus | Muridae | 29947 |
| MF618253 | Murine_hepatitis_virus_A59_MHV-ExoN(-)_P250 | Betacoronavirus | Muridae | 29893 |
| MF924725 | Infectious_bronchitis_virus_K2 | Gammacoronavirus | Phasianidae | 27626 |
| MG021194 | Infectious_bronchitis_virus_(gammaCoV)_AvCov_Ck_Italy_624I_96 | Gammacoronavirus | Phasianidae | 27552 |
| MG021451 | MERS-related_CoV_NL13845 | Betacoronavirus | Chiroptera | 30113 |
| MG021452 | MERS-related_CoV_NL140422 | Betacoronavirus | Chiroptera | 30113 |
| MG197727 | Infectious_bronchitis_virus_GD_QY16 | Gammacoronavirus | Phasianidae | 27696 |
| MG233398 | Infectious_bronchitis_virus_IS-1494 | Gammacoronavirus | Phasianidae | 27674 |
| MG334555 | Porcine_epidemic_diarrhea_virus_USA_OK10240-8 | Alphacoronavirus | Suidae | 27438 |
| MG428704 | Human_CoV_NL63_Kilifi_HH_5402 | Alphacoronavirus | Hominidae | 27832 |
| MG596802 | MERS-related_CoV_Bat-CoV_H.savii_Italy_206645 | Betacoronavirus | Chiroptera | 30048 |
| MG738154 | Infectious_bronchitis_virus_IBS037A | Gammacoronavirus | Phasianidae | 27669 |
| MG738155 | Infectious_bronchitis_virus_IBS130 | Gammacoronavirus | Phasianidae | 27660 |
| MG762674 | Rousettus_bat_CoV_HKU9_Rousettus_spp | Betacoronavirus | Chiroptera | 29134 |
| MG772933 | Bat_SARS-like_CoV_bat-SL-CoVZC45 | Betacoronavirus | Chiroptera | 29802 |
| MG772934 | Bat_SARS-like_CoV_bat-SL-CoVZXC21 | Betacoronavirus | Chiroptera | 29732 |
| MG812376 | Sparrow_deltaCoV_ISU690-7 | Deltacoronavirus | Passeridae | 25796 |
| MG812377 | Sparrow_deltaCoV_ISU42824 | Deltacoronavirus | Passeridae | 25710 |
| MG812378 | Sparrow_deltaCoV_ISU73347 | Deltacoronavirus | Passeridae | 25685 |
| MG893511 | Feline_CoV_Felix | Alphacoronavirus | Felidae | 29298 |
| MG913342 | Avian_CoV_(AvCoV)_Gallus_gallus_Brazil_sample_38_GI-11 | Gammacoronavirus | Phasianidae | 27618 |
| MG916903 | Bat_CoV_(BtCoV)_Rh_YN2012_(BtCoV)_Rs4259 | Unclassified_Coronavirinae | Chiroptera | 29076 |
| MG916904 | Bat_CoV_(BtCoV)_Rh_YN2012_(BtCoV)_Ra13591 | Unclassified_Coronavirinae | Chiroptera | 28751 |
| MG923471 | MERS-CoV_camel_Burkina_Faso_CIRAD-HKU785 | Betacoronavirus | Camelidae | 29721 |
| MG923472 | MERS-CoV_camel_Nigeria_NS004 | Betacoronavirus | Camelidae | 29680 |
| MG923475 | MERS-CoV_camel_Nigeria_NV1657 | Betacoronavirus | Camelidae | 29462 |
| MH002337 | Tylonycteris_bat_CoV_HKU4_CZ01 | Betacoronavirus | Chiroptera | 30255 |
| MH002342 | Pipistrellus_bat_CoV_HKU5_YD13403 | Betacoronavirus | Chiroptera | 30529 |
| MH021175 | Avian_CoV_D274 | Gammacoronavirus | Phasianidae | 27599 |
| MH181793 | Infectious_bronchitis_virus_HH06 | Gammacoronavirus | Phasianidae | 27672 |
| MH243319 | Porcine_epidemic_diarrhea_virus_KNU-1807 | Alphacoronavirus | Suidae | 28017 |
| MH532440 | Quail_deltaCoV_G032 | Deltacoronavirus | Phasianidae | 25881 |
| MH615810 | SADS-CoV-CH-FJWT | Alphacoronavirus | Suidae | 27184 |
| MH687938 | AlphaCoV_sp._VZ_16715_39_c1 | Alphacoronavirus | Chiroptera | 28349 |
| MH687941 | AlphaCoV_sp._VZ_16715_47_c1 | Alphacoronavirus | Chiroptera | 28008 |
| MH687951 | AlphaCoV_sp._VZ_16715_78 | Alphacoronavirus | Chiroptera | 28313 |
| MH687958 | AlphaCoV_sp._VZ_16845_87 | Alphacoronavirus | Chiroptera | 28036 |
| MH687960 | AlphaCoV_sp._VZ_17819_22 | Alphacoronavirus | Chiroptera | 27303 |
| MH687962 | AlphaCoV_sp._VZ_17819_50 | Alphacoronavirus | Chiroptera | 27752 |
| MH687966 | AlphaCoV_sp._VZ_20745_6 | Alphacoronavirus | Chiroptera | 28389 |
| MH687967 | AlphaCoV_sp._VZ_20745_8 | Alphacoronavirus | Chiroptera | 27726 |
| MH687968 | BetaCoV_sp._VZ_16715_52 | Betacoronavirus | Muridae | 31083 |
| MH687969 | BetaCoV_sp._VZ_20724_33 | Betacoronavirus | Muridae | 31352 |
| MH687970 | BetaCoV_sp._VZ_20724_34_c12 | Betacoronavirus | Muridae | 31038 |
| MH687971 | BetaCoV_sp._VZ_20724_34_c13 | Betacoronavirus | Muridae | 31171 |
| MH687977 | BetaCoV_sp._VZ_22084_10 | Betacoronavirus | Muridae | 31327 |
| MH708124 | Porcine_deltaCoV_HNZK-04 | Deltacoronavirus | Suidae | 25454 |
| MH779856 | Infectious_bronchitis_virus_ArkGA_P1 | Gammacoronavirus | Phasianidae | 27603 |
| MH817484 | Feline_1_(FCoV)-SB22 | Alphacoronavirus | Felidae | 29137 |
| MH878976 | Infectious_bronchitis_virus_VFAR-047 | Gammacoronavirus | Phasianidae | 27467 |
| MH924835 | Infectious_bronchitis_virus_(gammaCoV)_I0636_16 | Gammacoronavirus | Phasianidae | 27639 |
| MH938448 | AlphaCoV_Bat-CoV_P.kuhlii_Italy_206645-41 | Alphacoronavirus | Chiroptera | 27862 |
| MH938450 | AlphaCoV_Bat-CoV_P.kuhlii_Italy_206679-3 | Alphacoronavirus | Chiroptera | 28146 |
| MK032177 | Infectious_bronchitis_virus_(gammaCoV)_I0724_17 | Gammacoronavirus | Phasianidae | 27681 |
| MK062183 | SARS_CoV_Urbani_icSARS-C7 | Betacoronavirus | Hominidae | 29874 |
| MK129253 | MERS-CoV_KOR_KCDC_001_2018-TSVi | Betacoronavirus | Hominidae | 30150 |
| MK142676 | Infectious_bronchitis_virus_ahysx-1 | Gammacoronavirus | Phasianidae | 27718 |
| MK211369 | Coronavirus_BtSk-AlphaCoV_GX2018A | Alphacoronavirus | Chiroptera | 28303 |
| MK211372 | Coronavirus_BtSk-AlphaCoV_GX2018D | Alphacoronavirus | Chiroptera | 28238 |
| MK211373 | Coronavirus_BtRs-AlphaCoV_YN2018 | Alphacoronavirus | Chiroptera | 29109 |
| MK211374 | Coronavirus_BtRl-BetaCoV_SC2018 | Betacoronavirus | Chiroptera | 29648 |
| MK211378 | Coronavirus_BtRs-BetaCoV_YN2018D | Betacoronavirus | Chiroptera | 30213 |
| MK211379 | Coronavirus_BtRt-BetaCoV_GX2018 | Betacoronavirus | Chiroptera | 29752 |
| MK217372 | Infectious_bronchitis_virus_I0221_17 | Gammacoronavirus | Phasianidae | 27617 |
| MK217373 | Infectious_bronchitis_virus_I0725_17 | Gammacoronavirus | Phasianidae | 27708 |
| MK217375 | Infectious_bronchitis_virus_I1209_16 | Gammacoronavirus | Phasianidae | 27660 |
| MK330604 | Porcine_deltaCoVN_Sichuan | Deltacoronavirus | Suidae | 25402 |
| MK334045 | Human_CoV_NL63inaGD05 | Alphacoronavirus | Hominidae | 27544 |
| MK423876 | Pheasant_CoV_(gammaCoV)_phina_I0710_17 | Gammacoronavirus | Phasianidae | 27655 |
| MK462255 | MERS-related_CoV_Hu_Riyadh-KSA-18014504 | Betacoronavirus | Hominidae | 30052 |
| MK472067 | AlphaCoV_sp._WA1087 | Alphacoronavirus | Chiroptera | 28170 |
| MK472068 | AlphaCoV_sp._WA2028 | Alphacoronavirus | Chiroptera | 27435 |
| MK472070 | AlphaCoV_sp._WA3607 | Alphacoronavirus | Chiroptera | 28009 |
| MK472071 | AlphaCoV_sp._WAAlc1 | Alphacoronavirus | Chiroptera | 27535 |
| MK492263 | Bat_CoV_(BtCoV)92 | Unclassified_Coronavirinae | Chiroptera | 29585 |
| MK581201 | Infectious_bronchitis_virus_(gammaCoV)_79 | Gammacoronavirus | Phasianidae | 27640 |
| MK581202 | Infectious_bronchitis_virus_(gammaCoV)_80 | Gammacoronavirus | Phasianidae | 27875 |
| MK581203 | Infectious_bronchitis_virus_(gammaCoV)_162 | Gammacoronavirus | Phasianidae | 27644 |
| MK581204 | Infectious_bronchitis_virus_(gammaCoV)_255 | Gammacoronavirus | Phasianidae | 27651 |
| MK581205 | Infectious_bronchitis_virus_(gammaCoV)_548 | Gammacoronavirus | Phasianidae | 27844 |
| MK581206 | Infectious_bronchitis_virus_(gammaCoV)_195 | Gammacoronavirus | Phasianidae | 27433 |
| MK581207 | Infectious_bronchitis_virus_(gammaCoV)_G103 | Gammacoronavirus | Phasianidae | 27791 |
| MK679660 | Hedgehog_CoV_1 | Betacoronavirus | Erinaceidae | 30172 |
| MK720944 | Tylonycteris_bat_CoV_HKU33_GZ151867 | Alphacoronavirus | Chiroptera | 27636 |
| MK720945 | Rhinolophus_bat_CoV_HKU32_TLC26A | Alphacoronavirus | Chiroptera | 29201 |
| MK878536 | Infectious_bronchitis_virus_GA9977 | Gammacoronavirus | Phasianidae | 27672 |
| MK907286 | Erinaceus_hedgehog_CoV_HKU31_F6 | Betacoronavirus | Erinaceidae | 29951 |
| MK993519 | Porcine_deltaCoVN_Sichuan | Deltacoronavirus | Suidae | 25380 |
| MN065811 | Bat_alphaCoV_(BtCoV)_008_16_M.bra_FIN | Alphacoronavirus | Chiroptera | 28119 |
| MN096598 | Infectious_bronchitis_virus_160501 | Gammacoronavirus | Phasianidae | 27680 |
| MN128087 | Infectious_bronchitis_virus_TW2575_98vac | Gammacoronavirus | Phasianidae | 27755 |
| MN128088 | Infectious_bronchitis_virus_TW2296_95w | Gammacoronavirus | Phasianidae | 27733 |
| MN165107 | Feline_CoV_XXN | Alphacoronavirus | Felidae | 29286 |
| MN262644 | Avian_CoV_CV10 | Gammacoronavirus | Phasianidae | 27523 |
| MN306053 | Human_CoV_OC43_SC9430 | Betacoronavirus | Hominidae | 30818 |
| MN486588 | Porcine_epidemic_diarrhea_virus_Ah2016f2 | Alphacoronavirus | Suidae | 27933 |
| MN512434 | Infectious_bronchitis_virus_(IBV)_17-035614 | Gammacoronavirus | Phasianidae | 27710 |
| MN512436 | Infectious_bronchitis_virus_(IBV)_18-048192T | Gammacoronavirus | Phasianidae | 27638 |
| MN512437 | Infectious_bronchitis_virus_(IBV)_18-048430 | Gammacoronavirus | Phasianidae | 27684 |
| MN514965 | Dromedary_camel_CoV_HKU23_(DcCoV)-NV1092 | Betacoronavirus | Camelidae | 30799 |
| MN514966 | Dromedary_camel_CoV_HKU23_(DcCoV)-NV1097 | Betacoronavirus | Camelidae | 31075 |
| MN599049 | Infectious_bronchitis_virus_GA_1476 | Gammacoronavirus | Phasianidae | 27666 |
| MN611517 | Rousettus_aegyptiacus_bat_CoV_229E-related_5425 | Alphacoronavirus | Chiroptera | 27619 |
| MN611520 | Pipistrellus_abramus_bat_CoV_HKU5-related_BY140568 | Betacoronavirus | Chiroptera | 30511 |
| MN611521 | Scotophilus_kuhlii_bat_CoV_512-related_HK140714 | Alphacoronavirus | Chiroptera | 27933 |
| MN611522 | Rhinolophus_affinis_bat_CoV_HKU2-related_160660 | Alphacoronavirus | Chiroptera | 26956 |
| MN611523 | Hipposideros_pomona_bat_CoV_HKU10-related_160942 | Alphacoronavirus | Chiroptera | 28514 |
| MN611525 | Hipposideros_pomona_bat_CoVB25B0025 | Alphacoronavirus | Chiroptera | 28169 |
| MN690608 | Bottlenose_dolphin_CoV_37112-1 | Gammacoronavirus | Delphinidae | 31728 |
| MT040335 | Pangolin_CoV_PCoV_GX-P5L | Betacoronavirus | Manidae | 29806 |
| MT121215 | SARS-CoV-2_SH01_human_2020N | Betacoronavirus | Hominidae | 29945 |
| MT262993 | SARS-Cov-2_Manga1_human_2020_PAK | Betacoronavirus | Hominidae | 29836 |
| MT339040 | SARS-CoV-2_human_USA_AZ-ASU2923 | Betacoronavirus | Hominidae | 29807 |
| NC_002645 | Human_CoV_229E | Alphacoronavirus | Hominidae | 27317 |
| NC_006577 | Human_CoV_HKU1 | Betacoronavirus | Hominidae | 29926 |
| NC_009019 | Bat_CoV_HKU4-1 | Betacoronavirus | Chiroptera | 30286 |
| NC_009020 | Bat_CoV_HKU5-1 | Betacoronavirus | Chiroptera | 30482 |
| NC_009021 | Bat_CoV_HKU9-1 | Betacoronavirus | Chiroptera | 29114 |
| NC_010437 | Bat_CoV_1A | Alphacoronavirus | Chiroptera | 28326 |
| NC_010438 | Bat_CoV_HKU8 | Alphacoronavirus | Chiroptera | 28773 |
| NC_010646 | Beluga_Whale_CoV_SW1 | Gammacoronavirus | Cetacea | 31686 |
| NC_011547 | Bulbul_CoV_HKU11-934 | Deltacoronavirus | Pycnonotidae | 26487 |
| NC_011549 | Thrush_CoV_HKU12-600 | Deltacoronavirus | Turdidae | 26396 |
| NC_011550 | Munia_CoV_HKU13-3514 | Deltacoronavirus | Ploceidae | 26552 |
| NC_014470 | Bat_CoV_BM48-31_BGR | Betacoronavirus | Chiroptera | 29276 |
| NC_016991 | White-eye_CoV_HKU16 | Deltacoronavirus | Zosteropidae | 26041 |
| NC_016992 | Sparrow_CoV_HKU17 | Deltacoronavirus | Passeridae | 26083 |
| NC_016993 | Magpie-robin_CoV_HKU18 | Deltacoronavirus | Muscicapidae | 26689 |
| NC_016994 | Night-heron_CoV_HKU19 | Deltacoronavirus | Ardeidae | 26077 |
| NC_016995 | Wigeon_CoV_HKU20 | Deltacoronavirus | Anatidae | 26227 |
| NC_016996 | Common-moorhen_CoV_HKU21 | Deltacoronavirus | Rallidae | 26223 |
| NC_018871 | Rousettus_bat_CoV_HKU10 | Alphacoronavirus | Chiroptera | 28494 |
| NC_022103 | Bat_CoV_CDPHE15_USA | Alphacoronavirus | Chiroptera | 28035 |
| NC_023760 | Mink_CoV_WD1127 | Alphacoronavirus | Mustelidae | 28941 |
| NC_025217 | Bat_Hp-betaCoV_Zhejiang2013 | Betacoronavirus | Chiroptera | 31491 |
| NC_028806 | Swine_enteric_CoV_Italy_213306 | Alphacoronavirus | Suidae | 28111 |
| NC_028811 | BtMr-AlphaCoV_SAX2011 | Alphacoronavirus | Chiroptera | 27935 |
| NC_028814 | BtRf-AlphaCoV_HuB2013 | Alphacoronavirus | Chiroptera | 27608 |
| NC_028824 | BtRf-AlphaCoV_YN2012 | Alphacoronavirus | Chiroptera | 26975 |
| NC_028833 | BtNv-AlphaCoV_SC2013 | Alphacoronavirus | Chiroptera | 27783 |
| NC_030292 | Ferret_CoV_FRCoV-NL-2010 | Alphacoronavirus | Mustelidae | 28434 |
| NC_030886 | Rousettus_bat_CoV_GCCDC1_356 | Betacoronavirus | Chiroptera | 30161 |
| NC_032107 | NL63-related_bat_CoV_BtKYNL63-9a | Alphacoronavirus | Chiroptera | 28363 |
| NC_032730 | Lucheng_Rn_rat_CoV_Lucheng-19 | Alphacoronavirus | Muridae | 28763 |
| NC_034440 | Bat_CoV_PREDICT_PDF-2180 | Unclassified_Coronavirinae | Chiroptera | 29642 |
| NC_034972 | Coronavirus_AcCoV-JC34 | Alphacoronavirus | Muridae | 27682 |
| NC_039207 | ErinaceusCoV_2012-174_GER | Betacoronavirus | Erinaceidae | 30148 |
| NC_046964 | AlphaCoV_Bat-CoV_3398-19 | Alphacoronavirus | Chiroptera | 28128 |
| NC_046965 | Canada_goose_CoV_Cambridge_Bay | Gammacoronavirus | Anatidae | 28539 |

**Supplementary Table S2** The 26 coronavirus genomes added in the phylogenetic analysis for the subfamily Orthocoronavirinae

| **Accession ID** | **Strain** | **Genus** | **Host** | **Length (bp)** |
| --- | --- | --- | --- | --- |
| FJ959407 | Severe acute respiratory syndrome-related coronavirus | Betacoronavirus | Viverridae | 29694 |
| AY515512 | Severe acute respiratory syndrome-related coronavirus | Betacoronavirus | Viverridae | 29731 |
| AY572034 | Severe acute respiratory syndrome-related coronavirus | Betacoronavirus | Viverridae | 29540 |
| AY304486 | SARS coronavirus SZ3 | Betacoronavirus | Viverridae | 29741 |
| MN611518 | Miniopterus pusillus bat coronavirus HKU8-related | Alphacoronavirus | Vespertilionidae  (Chiroptera) | 28751 |
| MN611524 | Miniopterus schreibersii bat coronavirus 1-related | Alphacoronavirus | Vespertilionidae  (Chiroptera) | 28316 |
| MN611519 | Tylonycteris pachypus bat coronavirus HKU4-related | Betacoronavirus | Vespertilionidae  (Chiroptera) | 30224 |
| KY417142 | Severe acute respiratory syndrome-related coronavirus | Betacoronavirus | Vespertilionidae  (Chiroptera) | 29725 |
| FJ376620 | Bulbul coronavirus HKU11 | Deltacoronavirus | Pycnonotidae | 26476 |
| LC364343 | Houbara coronavirus UAE-HKU28 | Deltacoronavirus | Otididae | 26155 |
| MT072864 | Pangolin coronavirus | Betacoronavirus | Manidae | 29795 |
| MG518518 | Betacoronavirus 1 | Betacoronavirus | Cervidae | 31034 |
| FJ425187 | Betacoronavirus 1 | Betacoronavirus | Cervidae | 31020 |
| FJ425188 | Betacoronavirus 1 | Betacoronavirus | Cervidae | 30995 |
| MG977444 | Betacoronavirus 1 | Betacoronavirus | Hominidae | 30571 |
| KF906251 | Betacoronavirus 1 | Betacoronavirus | Camelidae | 31052 |
| JQ410000 | Human coronavirus 229E | Alphacoronavirus | Camelidae | 27374 |
| MN507638 | Middle East respiratory syndrome-related coronavirus | Betacoronavirus | Camelidae | 30033 |
| MH043953 | Betacoronavirus 1 | Betacoronavirus | Bovidae | 31032 |
| MH810163 | Betacoronavirus 1 | Betacoronavirus | Bovidae | 31032 |
| KU558922 | Betacoronavirus 1 | Betacoronavirus | Bovidae | 31039 |
| FJ425184 | Betacoronavirus 1 | Betacoronavirus | Bovidae | 30995 |
| EF424621 | Betacoronavirus 1 | Betacoronavirus | Bovidae | 30995 |
| MT457390 | SARS-CoV-2 (mink/NED/NB01_01KS/2020) | Betacoronavirus | Mustelidae | 29746 |
| MT365033 | SARS-CoV-2 (tiger/NY/040420/2020) | Betacoronavirus | Felidae | 29897 |
| MN996532 | Bat coronavirus RaTG13 | Betacoronavirus | Rhinolophidae (Chiroptera) | 29855 |

**Supplementary Table S3** The 269 coronavirus genomes used in the analysis of apparent horizontal gene transfer (HGT) events for the beta genus.

| **Accession ID** | **Strain** | **Genus** | **Host** | **Length (bp)** |
| --- | --- | --- | --- | --- |
| MH810163 | Yak_CoV_YAK_HY24 | Betacoronavirus | Bovidae | 31032 |
| FJ425187 | White-tailed_deer_CoV_US_OH-WD470 | Betacoronavirus | Cervidae | 31020 |
| MG518518 | Water_deer_CoV_W17-18 | Betacoronavirus | Cervidae | 31034 |
| MN611519 | Tylonycteris_pachypus_bat_CoV_HKU4-related_GZ131656 | Betacoronavirus | Chiroptera | 30224 |
| MH002337 | Tylonycteris_bat_CoV_HKU4_CZ01 | Betacoronavirus | Chiroptera | 30255 |
| KY352407 | Severe_acute_respiratory_syndrome-related_CoV_BtKY72 | Betacoronavirus | Chiroptera | 29274 |
| KU973692 | SARS-related_CoV_F46 | Betacoronavirus | Chiroptera | 29722 |
| MT365033 | SARS-CoV-2_tiger_NY_040420 | Betacoronavirus | Felidae | 29746 |
| MT457390 | SARS-CoV-2_mink_NED_NB01_01KS | Betacoronavirus | Mustelidae | 29897 |
| MK062183 | SARS_CoV_Urbani_icSARS-C7 | Betacoronavirus | Hominidae | 29874 |
| AY559083 | SARS_CoV_Sin3408 | Betacoronavirus | Hominidae | 29767 |
| AY463060 | SARS_CoV_ShanghaiQXC2 | Betacoronavirus | Hominidae | 29013 |
| AY515512 | SARS_CoV_HC_SZ_61_03 | Betacoronavirus | Viverridae | 29731 |
| AY572034 | SARS_CoV_civet007 | Betacoronavirus | Viverridae | 29540 |
| FJ959407 | SARS_CoV_A001 | Betacoronavirus | Viverridae | 29694 |
| MG762674 | Rousettus_bat_CoV_HKU9_Rousettus_spp | Betacoronavirus | Chiroptera | 29134 |
| NC_030886 | Rousettus_bat_CoV_GCCDC1_356 | Betacoronavirus | Chiroptera | 30161 |
| KF569996 | Rhinolophus_affinis_CoV_LYRa11 | Betacoronavirus | Chiroptera | 29805 |
| JF792616 | Rat_CoV_681 | Betacoronavirus | Muridae | 31286 |
| JN874560 | Rabbit_CoV_HKU14_HKU14-3 | Betacoronavirus | Leporidae | 31116 |
| JN874562 | Rabbit_CoV_HKU14_HKU14-10 | Betacoronavirus | Leporidae | 30904 |
| MH002342 | Pipistrellus_bat_CoV_HKU5_YD13403 | Betacoronavirus | Chiroptera | 30529 |
| MN611520 | Pipistrellus_abramus_bat_CoV_HKU5-related_BY140568 | Betacoronavirus | Chiroptera | 30511 |
| DQ011855 | PHEV_VW572 | Betacoronavirus | Suidae | 30480 |
| KY419112 | PHEV_CoV_USA-15TOSU1765 | Betacoronavirus | Suidae | 30531 |
| KY419109 | PHEV_CoV_USA-15TOSU1655 | Betacoronavirus | Suidae | 29906 |
| KY419110 | PHEV_CoV_USA-15TOSU1362 | Betacoronavirus | Suidae | 30515 |
| KY994645 | PHEV_CoV_JL | Betacoronavirus | Suidae | 30684 |
| MT040335 | Pangolin_CoV_PCoV_GX-P5L | Betacoronavirus | Manidae | 29806 |
| MT072864 | Pangolin_CoV_PCoV_GX-P2V | Betacoronavirus | Manidae | 29795 |
| JQ173883 | Murine_hepatitis_virus_S_3239-17 | Betacoronavirus | Muridae | 31161 |
| AB551247 | Murine_hepatitis_virus_MHV-MI | Betacoronavirus | Muridae | 31428 |
| AC_000192 | Murine_hepatitis_virus_JHM | Betacoronavirus | Muridae | 31526 |
| MF618252 | Murine_hepatitis_virus_A59_WT-MHV_P250 | Betacoronavirus | Muridae | 29947 |
| MF618253 | Murine_hepatitis_virus_A59_MHV-ExoN(-)_P250 | Betacoronavirus | Muridae | 29893 |
| AF201929 | Murine_hepatitis_virus_2 | Betacoronavirus | Muridae | 31276 |
| FJ647222 | Murine_CoV_SA59_RJHM | Betacoronavirus | Muridae | 31283 |
| FJ647224 | Murine_CoV_MHV-3 | Betacoronavirus | Muridae | 31448 |
| FJ647223 | Murine_CoV_MHV-1 | Betacoronavirus | Muridae | 31386 |
| JX169866 | Murine_CoV_JHM-WU-Dns2 | Betacoronavirus | Muridae | 30729 |
| JX169867 | Murine_CoV_JHM.WU | Betacoronavirus | Muridae | 31051 |
| KP887098 | Murine_CoV_AM2 | Betacoronavirus | Muridae | 31409 |
| MG021452 | MERS-related_CoV_NL140422 | Betacoronavirus | Chiroptera | 30113 |
| MG021451 | MERS-related_CoV_NL13845 | Betacoronavirus | Chiroptera | 30113 |
| MN507638 | MERS-related_CoV_llama-passaged-Qatar15 | Betacoronavirus | Camelidae | 30033 |
| MK462255 | MERS-related_CoV_Hu_Riyadh-KSA-18014504 | Betacoronavirus | Hominidae | 30052 |
| KY688119 | MERS-related_CoV_Hu_Aseer-KSA-Rs924 | Betacoronavirus | Hominidae | 30061 |
| MG596802 | MERS-related_CoV_Bat-CoV_H.savii_Italy_206645 | Betacoronavirus | Chiroptera | 30048 |
| MK129253 | MERS-CoV_KOR_KCDC_001_2018-TSVi | Betacoronavirus | Hominidae | 30150 |
| MG923475 | MERS-CoV_camel_Nigeria_NV1657 | Betacoronavirus | Camelidae | 29462 |
| MG923472 | MERS-CoV_camel_Nigeria_NS004 | Betacoronavirus | Camelidae | 29680 |
| MG923471 | MERS-CoV_camel_Burkina_Faso_CIRAD-HKU785 | Betacoronavirus | Camelidae | 29721 |
| KU710265 | MERS_CoV_Hu_Taif_KSA-7032_2014_S530del | Betacoronavirus | Hominidae | 29566 |
| KU851859 | MERS_CoV_Hu_Jeddah-KSA-3RS2702 | Betacoronavirus | Hominidae | 30076 |
| KX442564 | Hypsugo_bat_CoV_HKU25_YD131305 | Betacoronavirus | Chiroptera | 30498 |
| AY585229 | Human_CoV_OC43_serotype_OC43-Paris | Betacoronavirus | Hominidae | 30744 |
| MN306053 | Human_CoV_OC43_SC9430 | Betacoronavirus | Hominidae | 30818 |
| KP198610 | Human_CoV_OC43_2058A_10 | Betacoronavirus | Hominidae | 30752 |
| DQ415904 | Human_CoV_HKU1_N6_genotype_A | Betacoronavirus | Hominidae | 29295 |
| DQ415901 | Human_CoV_HKU1_N24_genotype_A | Betacoronavirus | Hominidae | 30097 |
| DQ415900 | Human_CoV_HKU1_N23_genotype_A | Betacoronavirus | Hominidae | 29812 |
| DQ415899 | Human_CoV_HKU1_N22_genotype_C | Betacoronavirus | Hominidae | 29905 |
| DQ415897 | Human_CoV_HKU1_N20_genotype_C | Betacoronavirus | Hominidae | 29845 |
| DQ415896 | Human_CoV_HKU1_N19_genotype_A | Betacoronavirus | Hominidae | 29860 |
| DQ415913 | Human_CoV_HKU1_N17_genotype_C | Betacoronavirus | Hominidae | 29785 |
| DQ415909 | Human_CoV_HKU1_N13_genotype_A | Betacoronavirus | Hominidae | 29767 |
| DQ415908 | Human_CoV_HKU1_N11_genotype_A | Betacoronavirus | Hominidae | 29947 |
| AY884001 | Human_CoV_HKU1_genotype_B | Betacoronavirus | Hominidae | 29815 |
| KT779555 | Human_CoV_HKU1_BJ01-p3 | Betacoronavirus | Hominidae | 29887 |
| NC_006577 | Human_CoV_HKU1 | Betacoronavirus | Hominidae | 29926 |
| KF686344 | HKU1_human_USA_HKU1-15 | Betacoronavirus | Hominidae | 29695 |
| KF686343 | HKU1_human_USA_HKU1-13 | Betacoronavirus | Hominidae | 29983 |
| MK679660 | Hedgehog_CoV_1 | Betacoronavirus | Erinaceidae | 30172 |
| EF424623 | Giraffe_CoV_US_OH3 | Betacoronavirus | Giraffidae | 31002 |
| NC_039207 | ErinaceusCoV_2012-174_GER | Betacoronavirus | Erinaceidae | 30148 |
| MK907286 | Erinaceus_hedgehog_CoV_HKU31_F6 | Betacoronavirus | Erinaceidae | 29951 |
| KC545386 | Erinaceus_CoV_2012-216_GER | Betacoronavirus | Erinaceidae | 30175 |
| LC061272 | Equine_CoV_Tokachi09 | Betacoronavirus | Equidae | 30782 |
| LC061273 | Equine_CoV_Obihiro12-1 | Betacoronavirus | Equidae | 30916 |
| EF446615 | Equine_CoV_NC99 | Betacoronavirus | Equidae | 30992 |
| MN514966 | Dromedary_camel_CoV_HKU23_(DcCoV)-NV1097 | Betacoronavirus | Camelidae | 31075 |
| MN514965 | Dromedary_camel_CoV_HKU23_(DcCoV)-NV1092 | Betacoronavirus | Camelidae | 30799 |
| KC869678 | Coronavirus_Neoromicia_PML-PHE1_RSA | Betacoronavirus | Chiroptera | 30111 |
| MK211379 | Coronavirus_BtRt-BetaCoV_GX2018 | Betacoronavirus | Chiroptera | 29752 |
| MK211378 | Coronavirus_BtRs-BetaCoV_YN2018D | Betacoronavirus | Chiroptera | 30213 |
| MK211374 | Coronavirus_BtRl-BetaCoV_SC2018 | Betacoronavirus | Chiroptera | 29648 |
| KT368891 | Camel_CoV_HKU23_camel | Betacoronavirus | Camelidae | 31041 |
| KJ473821 | BtVs-BetaCoV_SC2013 | Betacoronavirus | Chiroptera | 30423 |
| KJ473814 | BtRs-BetaCoV_HuB2013 | Betacoronavirus | Chiroptera | 29658 |
| KJ473815 | BtRs-BetaCoV_GX2013 | Betacoronavirus | Chiroptera | 29161 |
| KJ473811 | BtRf-BetaCoV_JL2012 | Betacoronavirus | Chiroptera | 29037 |
| AF220295 | Bovine_CoV_Quebec | Betacoronavirus | Bovidae | 31100 |
| MH043953 | Bovine_CoV_4-17-25 | Betacoronavirus | Bovidae | 31032 |
| MH687977 | BetaCoV_sp._VZ_22084_10 | Betacoronavirus | Muridae | 31327 |
| MH687971 | BetaCoV_sp._VZ_20724_34_c13 | Betacoronavirus | Muridae | 31171 |
| MH687970 | BetaCoV_sp._VZ_20724_34_c12 | Betacoronavirus | Muridae | 31038 |
| MH687969 | BetaCoV_sp._VZ_20724_33 | Betacoronavirus | Muridae | 31352 |
| MH687968 | BetaCoV_sp._VZ_16715_52 | Betacoronavirus | Muridae | 31083 |
| KU558922 | BetaCoV_1_Buffalo_CoV_B1-24F | Betacoronavirus | Bovidae | 31039 |
| KP886808 | Bat_SARS-like_CoV_YNLF_31C | Betacoronavirus | Chiroptera | 29723 |
| KY417150 | Bat_SARS-like_CoV_Rs4874 | Betacoronavirus | Chiroptera | 30311 |
| KY417148 | Bat_SARS-like_CoV_Rs4247 | Betacoronavirus | Chiroptera | 29743 |
| KC881006 | Bat_SARS-like_CoV_Rs3367 | Betacoronavirus | Chiroptera | 29792 |
| KY417145 | Bat_SARS-like_CoV_Rf4092 | Betacoronavirus | Chiroptera | 29710 |
| MG772934 | Bat_SARS-like_CoV_bat-SL-CoVZXC21 | Betacoronavirus | Chiroptera | 29732 |
| MG772933 | Bat_SARS-like_CoV_bat-SL-CoVZC45 | Betacoronavirus | Chiroptera | 29802 |
| KY417142 | Bat_SARS-like_CoV_As6526 | Betacoronavirus | Chiroptera | 29725 |
| FJ588686 | Bat_SARS_CoV_Rs672 | Betacoronavirus | Chiroptera | 29059 |
| DQ412043 | Bat_SARS_CoV_Rm1 | Betacoronavirus | Chiroptera | 29749 |
| DQ412042 | Bat_SARS_CoV_Rf1 | Betacoronavirus | Chiroptera | 29709 |
| GQ153543 | Bat_SARS_CoV_HKU3-8 | Betacoronavirus | Chiroptera | 29681 |
| GQ153548 | Bat_SARS_CoV_HKU3-13 | Betacoronavirus | Chiroptera | 29677 |
| DQ022305 | Bat_SARS_CoV_HKU3-1 | Betacoronavirus | Chiroptera | 29728 |
| NC_025217 | Bat_Hp-betaCoV_Zhejiang2013 | Betacoronavirus | Chiroptera | 31491 |
| JX993987 | Bat_CoV_Rp_Shaanxi2011 | Betacoronavirus | Chiroptera | 29484 |
| HM211098 | Bat_CoV_HKU9-5-1 | Betacoronavirus | Chiroptera | 29136 |
| EF065516 | Bat_CoV_HKU9-4 | Betacoronavirus | Chiroptera | 29155 |
| EF065514 | Bat_CoV_HKU9-2 | Betacoronavirus | Chiroptera | 29107 |
| HM211101 | Bat_CoV_HKU9-10-2 | Betacoronavirus | Chiroptera | 29122 |
| NC_009021 | Bat_CoV_HKU9-1 | Betacoronavirus | Chiroptera | 29114 |
| NC_009020 | Bat_CoV_HKU5-1 | Betacoronavirus | Chiroptera | 30482 |
| EF065508 | Bat_CoV_HKU4-4 | Betacoronavirus | Chiroptera | 30316 |
| NC_009019 | Bat_CoV_HKU4-1 | Betacoronavirus | Chiroptera | 30286 |
| JX993988 | Bat_CoV_Cp_Yunnan2011 | Betacoronavirus | Chiroptera | 29452 |
| NC_014470 | Bat_CoV_BM48-31_BGR | Betacoronavirus | Chiroptera | 29276 |
| MN996532 | Bat_coronavirus_RaTG13 | Betacoronavirus | Chiroptera | 29855 |
| DQ071615 | Bat SARS CoV Rp3/2004 | Betacoronavirus | Chiroptera | 29736 |
| DQ915164 | Bovine coronavirus isolate Alpaca | Betacoronavirus | Camelidae | 31076 |
| EF065510 | Bat coronavirus HKU5-2 | Betacoronavirus | Chiroptera | 30488 |
| EF424615 | Bovine coronavirus E-AH65 | Betacoronavirus | Bovidae | 31017 |
| EF424617 | Bovine coronavirus R-AH65 | Betacoronavirus | Bovidae | 31016 |
| EF424619 | Bovine coronavirus E-AH187 | Betacoronavirus | Bovidae | 30995 |
| EPI_ISL_402121 | SARS-CoV-2-IVDC-HB-05 | Betacoronavirus | Hominidae | 30637 |
| EPI_ISL_402123 | SARS-CoV-2-WH-01 | Betacoronavirus | Hominidae | 30645 |
| EPI_ISL_402124 | SARS-CoV-2-WIV04 | Betacoronavirus | Hominidae | 30637 |
| FJ415324 | Human enteric coronavirus 4408 | Betacoronavirus | Hominidae | 31029 |
| FJ425189 | Sambar deer coronavirus US/OH-WD388/1994 | Betacoronavirus | Cervidae | 30997 |
| FJ647220 | Murine coronavirus RA59/SJHM | Betacoronavirus | Muridae | 31429 |
| FJ647221 | Murine coronavirus repA59/RJHM | Betacoronavirus | Muridae | 31456 |
| FJ647226 | Murine coronavirus MHV-JHM.IA | Betacoronavirus | Muridae | 31473 |
| FJ647227 | Murine coronavirus repJHM/RA59 | Betacoronavirus | Muridae | 31250 |
| FJ938063 | Bovine coronavirus E-DB2-TC | Betacoronavirus | Bovidae | 31024 |
| FJ938064 | Bovine coronavirus E-AH187-TC | Betacoronavirus | Bovidae | 30995 |
| FJ938066 | Bovine respiratory coronavirus bovine/US/OH-440-TC/1996 | Betacoronavirus | Bovidae | 30953 |
| FJ938067 | Human enteric coronavirus strain 4408 | Betacoronavirus | Hominidae | 30953 |
| HM034837 | Human coronavirus HKU1 | Betacoronavirus | Hominidae | 29862 |
| HM211099 | Bat coronavirus HKU9-5-2 | Betacoronavirus | Chiroptera | 29112 |
| JF792617 | Rat coronavirus | Betacoronavirus | Muridae | 31274 |
| JN874561 | Rabbit coronavirus HKU14 | Betacoronavirus | Leporidae | 31096 |
| JX860640 | Canine respiratory coronavirus | Betacoronavirus | Canidae | 31028 |
| KC881005 | Bat SARS-like coronavirus RsSHC014 | Betacoronavirus | Chiroptera | 29787 |
| KF367457 | Bat SARS-like coronavirus WIV1 | Betacoronavirus | Chiroptera | 30309 |
| KF430201 | Human coronavirus HKU1 | Betacoronavirus | Hominidae | 29934 |
| KF530060 | Human coronavirus OC43 | Betacoronavirus | Hominidae | 30577 |
| KF530064 | Human coronavirus OC43 | Betacoronavirus | Hominidae | 30577 |
| KF530069 | Human coronavirus OC43 | Betacoronavirus | Hominidae | 30578 |
| KF530070 | Human coronavirus OC43 | Betacoronavirus | Hominidae | 30573 |
| KF530074 | Human coronavirus OC43 | Betacoronavirus | Hominidae | 30590 |
| KF530089 | Human coronavirus OC43 | Betacoronavirus | Hominidae | 30611 |
| KF530098 | Human coronavirus OC43 | Betacoronavirus | Hominidae | 30578 |
| KF600647 | Middle East respiratory syndrome-related coronavirus | Betacoronavirus | Hominidae | 30115 |
| KF686341 | Human coronavirus HKU1 | Betacoronavirus | Hominidae | 29832 |
| KF686342 | Human coronavirus HKU1 | Betacoronavirus | Hominidae | 29742 |
| KF906249 | Dromedary camel coronavirus HKU23 | Betacoronavirus | Camelidae | 31052 |
| KF923886 | Human coronavirus OC43 | Betacoronavirus | Hominidae | 30719 |
| KF923887 | Human coronavirus OC43 | Betacoronavirus | Hominidae | 30719 |
| KF923889 | Human coronavirus OC43 | Betacoronavirus | Hominidae | 30719 |
| KF923893 | Human coronavirus OC43 | Betacoronavirus | Hominidae | 30713 |
| KF923895 | Human coronavirus OC43 | Betacoronavirus | Hominidae | 30731 |
| KF923896 | Human coronavirus OC43 | Betacoronavirus | Hominidae | 30737 |
| KF923898 | Human coronavirus OC43 | Betacoronavirus | Hominidae | 30716 |
| KF923899 | Human coronavirus OC43 | Betacoronavirus | Hominidae | 30722 |
| KF923906 | Human coronavirus OC43 | Betacoronavirus | Hominidae | 30737 |
| KF923923 | Human coronavirus OC43 | Betacoronavirus | Hominidae | 30713 |
| KJ361501 | Middle East respiratory syndrome-related coronavirus | Betacoronavirus | Hominidae | 29880 |
| KJ361502 | Middle East respiratory syndrome-related coronavirus | Betacoronavirus | Hominidae | 29783 |
| KJ473812 | BtRf-BetaCoV/HeB2013 | Betacoronavirus | Chiroptera | 29443 |
| KJ473813 | BtRf-BetaCoV/SX2013 | Betacoronavirus | Chiroptera | 29461 |
| KJ473816 | BtRs-BetaCoV/YN2013 | Betacoronavirus | Chiroptera | 29142 |
| KJ473820 | BtPa-BetaCoV/GD2013 | Betacoronavirus | Chiroptera | 30480 |
| KJ473822 | BtTp-BetaCoV/GX2012 | Betacoronavirus | Chiroptera | 30247 |
| KJ477102 | Middle East respiratory syndrome-related coronavirus | Betacoronavirus | Camelidae | 29908 |
| KJ813439 | Middle East respiratory syndrome-related coronavirus | Betacoronavirus | Hominidae | 30123 |
| KJ958218 | Human coronavirus OC43 | Betacoronavirus | Hominidae | 30714 |
| KP198611 | Human coronavirus OC43 | Betacoronavirus | Hominidae | 30752 |
| KT368824 | Middle East respiratory syndrome-related coronavirus | Betacoronavirus | Camelidae | 30115 |
| KT368890 | Middle East respiratory syndrome-related coronavirus | Betacoronavirus | Camelidae | 30120 |
| KU131570 | Human coronavirus OC43 | Betacoronavirus | Hominidae | 30746 |
| KU558923 | Betacoronavirus 1 | Betacoronavirus | Bovidae | 30985 |
| KU886219 | Bovine coronavirus | Betacoronavirus | Bovidae | 30975 |
| KX108943 | Middle East respiratory syndrome-related coronavirus | Betacoronavirus | Camelidae | 30088 |
| KX344031 | Human coronavirus OC43 | Betacoronavirus | Hominidae | 30713 |
| KX432213 | Canine respiratory coronavirus | Betacoronavirus | Canidae | 30868 |
| KX442565 | Hypsugo bat coronavirus HKU25 | Betacoronavirus | Chiroptera | 30497 |
| KX538965 | Human coronavirus OC43 | Betacoronavirus | Hominidae | 30716 |
| KX538979 | Human coronavirus OC43 | Betacoronavirus | Hominidae | 30713 |
| KX982264 | Bovine coronavirus | Betacoronavirus | Bovidae | 30847 |
| KY014281 | Human coronavirus OC43 | Betacoronavirus | Hominidae | 30602 |
| KY369907 | Human coronavirus OC43 | Betacoronavirus | Hominidae | 30725 |
| KY417143 | Bat SARS-like coronavirus | Betacoronavirus | Chiroptera | 29741 |
| KY417144 | Bat SARS-like coronavirus | Betacoronavirus | Chiroptera | 29770 |
| KY417146 | Bat SARS-like coronavirus | Betacoronavirus | Chiroptera | 29782 |
| KY417147 | Bat SARS-like coronavirus | Betacoronavirus | Chiroptera | 29741 |
| KY417149 | Bat SARS-like coronavirus | Betacoronavirus | Chiroptera | 29743 |
| KY417151 | Bat SARS-like coronavirus | Betacoronavirus | Chiroptera | 30307 |
| KY417152 | Bat SARS-like coronavirus | Betacoronavirus | Chiroptera | 29769 |
| KY419103 | Porcine hemagglutinating encephalomyelitis virus | Betacoronavirus | Suidae | 30632 |
| KY673148 | Middle East respiratory syndrome-related coronavirus | Betacoronavirus | Hominidae | 30123 |
| KY674921 | Human coronavirus HKU1 | Betacoronavirus | Hominidae | 29820 |
| KY674943 | Human coronavirus HKU1 | Betacoronavirus | Hominidae | 29908 |
| KY967356 | Human coronavirus OC43 | Betacoronavirus | Hominidae | 30648 |
| MF000458 | Middle East respiratory syndrome-related coronavirus | Betacoronavirus | Hominidae | 30045 |
| MF314143 | Human coronavirus OC43 | Betacoronavirus | Hominidae | 30737 |
| MF374983 | Human coronavirus OC43 | Betacoronavirus | Hominidae | 30721 |
| MF374984 | Human coronavirus OC43 | Betacoronavirus | Hominidae | 30713 |
| MF593268 | Middle East respiratory syndrome-related coronavirus | Betacoronavirus | Chiroptera | 30009 |
| MF598596 | Middle East respiratory syndrome-related coronavirus | Betacoronavirus | Camelidae | 30123 |
| MF598616 | Middle East respiratory syndrome-related coronavirus | Betacoronavirus | Camelidae | 30123 |
| MF598624 | Middle East respiratory syndrome-related coronavirus | Betacoronavirus | Camelidae | 30123 |
| MF598631 | Middle East respiratory syndrome-related coronavirus | Betacoronavirus | Camelidae | 30123 |
| MF598638 | Middle East respiratory syndrome-related coronavirus | Betacoronavirus | Camelidae | 30123 |
| MG011342 | Middle East respiratory syndrome-related coronavirus | Betacoronavirus | Hominidae | 30096 |
| MG011356 | Middle East respiratory syndrome-related coronavirus | Betacoronavirus | Hominidae | 30096 |
| MG596803 | Middle East respiratory syndrome-related coronavirus | Betacoronavirus | Chiroptera | 30039 |
| MG757138 | Bovine coronavirus | Betacoronavirus | Bovidae | 31026 |
| MG757139 | Bovine coronavirus | Betacoronavirus | Bovidae | 31028 |
| MG912595 | Middle East respiratory syndrome-related coronavirus | Betacoronavirus | Hominidae | 30123 |
| MG923468 | Middle East respiratory syndrome-related coronavirus | Betacoronavirus | Camelidae | 30091 |
| MG923469 | Middle East respiratory syndrome-related coronavirus | Betacoronavirus | Camelidae | 29421 |
| MG923473 | Middle East respiratory syndrome-related coronavirus | Betacoronavirus | Camelidae | 29374 |
| MG977451 | Human coronavirus OC43 | Betacoronavirus | Hominidae | 30722 |
| MG977452 | Human coronavirus OC43 | Betacoronavirus | Hominidae | 30704 |
| MH013216 | Middle East respiratory syndrome-related coronavirus | Betacoronavirus | Hominidae | 30118 |
| MH029552 | Middle East respiratory syndrome-related coronavirus | Betacoronavirus | Hominidae | 30123 |
| MH043952 | Bovine coronavirus | Betacoronavirus | Bovidae | 31019 |
| MH043954 | Bovine coronavirus | Betacoronavirus | Bovidae | 31016 |
| MH043955 | Bovine coronavirus | Betacoronavirus | Bovidae | 31032 |
| MH310909 | Middle East respiratory syndrome-related coronavirus | Betacoronavirus | Hominidae | 30065 |
| MH310912 | Middle East respiratory syndrome-related coronavirus | Betacoronavirus | Hominidae | 30085 |
| MH432120 | Middle East respiratory syndrome-related coronavirus | Betacoronavirus | Hominidae | 30119 |
| MH454272 | Middle East respiratory syndrome-related coronavirus | Betacoronavirus | Hominidae | 30119 |
| MH687973 | Betacoronavirus sp. | Betacoronavirus | Muridae | 31323 |
| MH687974 | Betacoronavirus sp. | Betacoronavirus | Muridae | 31223 |
| MH687976 | Betacoronavirus sp. | Betacoronavirus | Muridae | 31095 |
| MH734114 | Middle East respiratory syndrome-related coronavirus | Betacoronavirus | Camelidae | 30033 |
| MK039552 | Middle East respiratory syndrome-related coronavirus | Betacoronavirus | Hominidae | 30123 |
| MK062181 | SARS coronavirus Urbani | Betacoronavirus | Hominidae | 29727 |
| MK167038 | Human coronavirus HKU1 | Betacoronavirus | Hominidae | 29921 |
| MK211375 | Coronavirus BtRs-BetaCoV/YN2018A | Betacoronavirus | Chiroptera | 29698 |
| MK211376 | Coronavirus BtRs-BetaCoV/YN2018B | Betacoronavirus | Chiroptera | 30256 |
| MK211377 | Coronavirus BtRs-BetaCoV/YN2018C | Betacoronavirus | Chiroptera | 29689 |
| MK462247 | Middle East respiratory syndrome-related coronavirus | Betacoronavirus | Hominidae | 30118 |
| MK564474 | Middle East respiratory syndrome-related coronavirus | Betacoronavirus | Camelidae | 30085 |
| MK967708 | Middle East respiratory syndrome-related coronavirus | Betacoronavirus | Camelidae | 30106 |
| MN026164 | Human coronavirus OC43 | Betacoronavirus | Hominidae | 30777 |
| MN306036 | Human coronavirus OC43 | Betacoronavirus | Hominidae | 30681 |
| MN306043 | Human coronavirus OC43 | Betacoronavirus | Hominidae | 30716 |
| MN310476 | Human coronavirus OC43 | Betacoronavirus | Hominidae | 30562 |
| MN310478 | Human coronavirus OC43 | Betacoronavirus | Hominidae | 30681 |
| MN514962 | Dromedary camel coronavirus HKU23 | Betacoronavirus | Camelidae | 31021 |
| MN514963 | Dromedary camel coronavirus HKU23 | Betacoronavirus | Camelidae | 31062 |
| MN723544 | Middle East respiratory syndrome-related coronavirus | Betacoronavirus | Hominidae | 30123 |
| NC_003045 | Bovine_coronavirus | Betacoronavirus | Unknown | 31028 |
| NC_012936 | Rat coronavirus Parker | Betacoronavirus | Muridae | 31250 |
| NC_017083 | Rabbit coronavirus HKU14 | Betacoronavirus | Leporidae | 31100 |
| NC_019843 | Middle East respiratory syndrome-related coronavirus | Betacoronavirus | Hominidae | 30119 |
| NC_026011 | Betacoronavirus HKU24 | Betacoronavirus | Muridae | 31249 |
| NC_045512 | Severe acute respiratory syndrome coronavirus 2 | Betacoronavirus | Hominidae | 29903 |
| U00735 | Bovine coronavirus | Betacoronavirus | Unknown | 31032 |

**Supplementary Figure S1** Rank abundance plot of top 20 protein clusters among alpha-coronavirus, beta-coronavirus, gamma-coronavirus and delta-coronavirus. Protein clusters with abundancy of more than 30% in a genus was selected to plot. Average copy number for each marker is shown as dots.

 
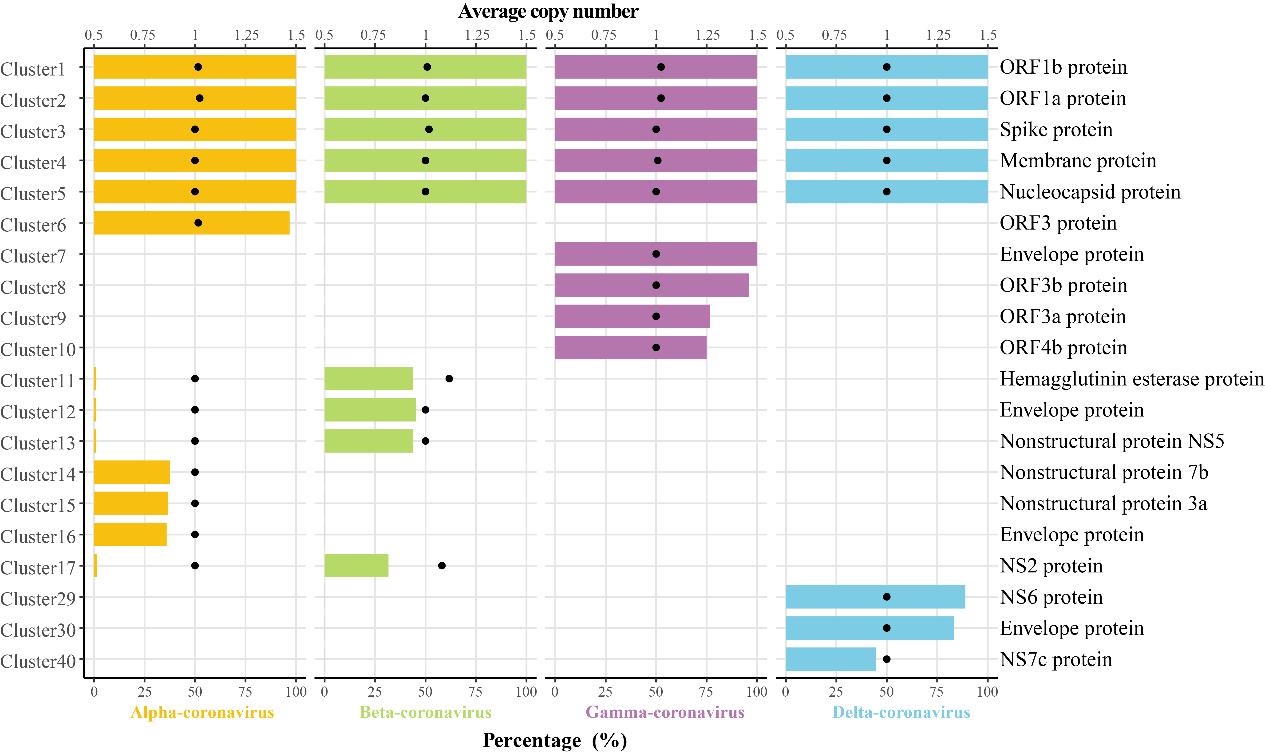
